# Multivalent Lipid MVL5 Micellar Nanoparticles Exhibit Dramatically Increased Loading of Paclitaxel with PEGylation Enhancing Human Cancer Cell Penetration Depth and Cytotoxicity

**DOI:** 10.1101/2025.05.24.655953

**Authors:** William S. Fisher, Aria Ghasemizadeh, Sherwin Roshan, Anna Goldstein, Jessica Douglas, Ramon Perez, Youli Li, Kai K. Ewert, Cyrus R. Safinya

## Abstract

Cationic liposomes (CLs) with chain-melted fluid membranes are promising nanocarriers of hydrophobic drugs in cancer chemotherapy, including the frequently employed drug paclitaxel (PTX). CL formulations containing univalent *N*-[2,3-dioleoyloxy-1-propyl]trimethylammonium chloride (DOTAP), like EndoTAG-1^TM^, have shown limited success in clinical trials and challenges like endosomal entrapment, limited PTX membrane solubility, and difficulty with *in vivo* tumor targeting remain. Incorporation of 10 mol% cone-shaped poly(ethylene glycol)-lipid (PEG-lipid) to DOTAP-containing CLs transitions a fraction of the particles to micellar nanodiscs. These PTX—loaded PEGylated CLs and nanodiscs show enhanced cellular uptake *in vitro* and improved tumor penetration and proapoptotic activity compared to bare CLs in an *in vivo* solid breast cancer tumor model. Formulations incorporating the multivalent cationic lipid MVL5 (+5e) at 50 mol% form nanoparticles (NPs) comprised almost entirely of nanodiscs, and transition at 75 mol% MVL5 to short micellar rods coexisting with spheres, with rods further transitioning to long flexible rods upon PEGylation. Here, we report on the finding that MVL5-based micellar NPs with disc, rod, and spherical morphologies dramatically improve the solubility of PTX in their fluid lipid membranes by nearly three-fold compared to reference CLs modeled on the EndoTAG-1^TM^ formulation. Cell viability assays revealed that this improved PTX solubility for MVL5 micellar NPs leads to improved cytotoxic efficacy, which is further improved by PEGylation. Remarkably, using fluorescent microscopy and image particle analysis, we find that the cellular uptake and penetration depth of MVL5 nanoparticles is significantly improved by PEGylation. The findings are consistent with a model where the rate-limiting step of PTX delivery by cationic lipid NPs is diffusion of endocytic vesicles containing NPs through the actin mesh near the cell surface combined with the hoping rate of PTX from endosomal membrane to nearby microtubules. PEGylated cationic lipid nanoparticles containing MVL5 therefore represent a very promising hydrophobic cancer drug delivery vehicle for nanomedicine applications.

## INTRODUCTION

Numerous therapeutically valuable chemotherapeutic drugs are hydrophobic and require carriers to solubilize them in aqueous environments and enable their delivery in the body. Paclitaxel (PTX) is a prominent example of such a drug and among the most common treatments for ovarian, breast, lung, and pancreatic cancers.^1–4^ PTX inhibits mitosis and induces cell death in cancer cells by stabilizing microtubules upon binding a specific hydrophobic pocket on β-tubulin.^3^ This requires that PTX remain solubilized in the body, as its insoluble, crystallized form is therapeutically inert. Common carriers of PTX used to achieve solubility in the body today include Taxol®, which uses the Kolliphor EL surfactant, and Abraxane, where PTX is bound to Albumin and complexed as Albumin nanoparticles.^5–8^ These are commonly associated with dose limiting toxicities from either the vehicle or the indiscriminate action of PTX against healthy cells throughout the body.^9–11^ To improve therapies using PTX, delivery vehicles for PTX that avoid the use of Kolliphor EL and its associated hypersensitivity reactions while increasing the specificity of PTX delivery to the tumor tissue are needed.

Liposomes are widely used carriers for both hydrophobic and hydrophilic small molecule drugs and biological (proteins and nucleic acid) therapeutics, owing to their relatively high biocompatibility and versatile cargo loading via their aqueous core or membrane.^5, 12, 13^ Incorporating PTX in the hydrophobic membrane of liposomes reduces toxicity compared to Taxol® and may enhance the maximum tolerable dose and biodistribution of PTX.^14–17^ This has led to the clinical deployment of several PTX-loaded liposomes. Lipusu®, whose composition is not publicly available, is approved for use in China.^18, 19^ LEP-ETU, which consists of the neutral lipid DOPC (1,2- dioleoyl-sn-glycero-3-phosphatidylcholine), cholesterol, the anionic lipid cardiolipin (90:5:5 mol ratio), and 3 mol% PTXL, is in phase II clinical trials in the United States.^20^ Finally, the cationic liposome (CL) formulation EndoTAG-1^TM^, which consists of the univalent cationic lipid DOTAP (2,3-dioleyloxypropyltrimethylammonium chloride), DOPC, and PTX at a 50:47:3 mol ratio, is in Phase III clinical trials in Taiwan.^21^ These liposomal PTX carriers all leverage the leaky neoangiogenic vasculature and defective lymphatic drainage around solid tumors to preferentially accumulate PTX at tumor tissue, the enhanced permeability and retention effect (EPR).^21–23^ CLs complement this passive targeting with an electrostatic attraction between them and the anionic sulfated proteoglycan layer that is observed to be enriched in neoangiogenic vasculature, creating an active targeting effect.^24–27^ We investigated cationic lipid nanoparticles because of their active targeting ability and the clinical success of CLs containing DOTAP so far, which serve as a benchmark for this study.

There remains significant room for optimization of CL carriers of PTX through improved drug loading capacity, biodistribution, and cellular uptake. For example, the efficacy of PTX-loaded CLs is determined in part by the solubility of PTX in their membrane, because the separation of PTX from the liposome membrane and formation of insoluble PTX crystals results in reduced bioavailability and, therefore, reduced cytotoxic efficacy.^21, 28–31^ Prior work shows that PTX membrane solubility at PTX contents at 3 mol%, which are used in liposomal formulations currently under clinical investigation, is limited and that this may negatively impact their therapeutic efficacy.^28^ Altered lipid structure can improve PTX solubility, as prior investigation of lipid tail structure revealed that double unsaturated tails significantly improved the solubility limit of PTX in CLs.^32^ Additionally, PTX-loaded CLs must circulate through the body and reach the tumor site to achieve their therapeutic effect. PEGylation, or the surface decoration of CLs with polyethylene glycol (PEG) using PEG-conjugated lipids, stabilizes CLs to prevent aggregation through non-specific interactions and is shown to extend their circulation time by delaying immune recognition and clearance.^13, 33^ Despite its importance for optimum biodistribution, EndoTAG-1^TM^ lacks PEGylation.

Beyond steric stabilization, the incorporation of 10 mol% PEG-lipid into CLs with fluid phase membranes above a critical concentration associated with the mushroom-brush transition prevents the formation of vesicles larger than 100 nm and drives the formation of distinct 10 to 20 nm disc-shaped micelles coexisting with vesicles, as revealed by cryogenic TEM (Cryo-TEM, see Supplemental Figure 1 in the Supplemental Information).^34, 35^ This is due to the positive spontaneous curvature (C_0_ > 0) of the cone-shaped PEG-lipids compared to the neutral spontaneous curvature (C_0_ = 0) of cylindrical DOPC and DOTAP lipids. When PEG lipids are incorporated at low molar concentrations (i.e. below 5 mol%), the adjacent PEG molecules that create their positive curvature are in a randomly packed, or mushroom, conformation and membranes can adopt a flat curvature. Above a critical molar concentration, PEG molecules in the mushroom conformation overlap and steric repulsion leads to the segregation of PEG molecules to high curvature regions of discoidal micelles, which reduces the curvature elastic energy cost of edge formation.^36, 37^ The altered shape and size of CLs and disc-micelles with 10 mol% PEG may confer advantages beyond steric stabilization. These include the finding that lipid nanoparticle morphology (disc versus sphere) affect their cell uptake through distinct endocytic pathways^38^ and, an increased number of collisions with endothelial cells in blood vessels, which is observed for non-spherical nanoparticles.^39^ Notably, incorporation of 10 mol% PEG in CLs significantly improves cellular uptake and cytotoxic efficacy compared to bare CLs *in vitro* and improves circulation time, extravasation, tumor accumulation, and apoptotic signal activation compared to bare and 5 mol% PEG-containing CLs, *in vivo*.^35, 40^

Lipids with highly charged multivalent headgroups are also expected to produce lipid assemblies with C_0_ > 0 and a recent Cryo-TEM study of cationic lipid nanoparticles (CLNPs) containing 50 mol% MVL5 (+5e, Figure 1) with DOPC as the remaining content, revealed that they form populations of almost entirely disc micelles (Supplemental Figure 2 in Supplemental Information)^34^ in contrast to the small fraction of disc nanoparticles formed with DOTAP/DOPC formulations containing 10 mol% PEG-lipid.^34, 35^ The multivalent cationic lipid MVL5 achieves a very high positive curvature as a function of the effective size of its headgroup,^41^ which results from the electrostatic repulsion created by its five ionizable amine groups that are positively charged at physiological pH. MVL5 is commercially available and has been studied in lipid only formulations and those complexed with nucleic acids (DNA, siRNA).^42–46^ The incorporation of MVL5 at 75 mol% further alters the structure of these particles, creating populations comprised of spherical micelles and short micellar rods (Supplemental Figure 2 in Supplemental Information) while the addition of 10 mol% PEG-lipid to 50 mol% MVL5 particles creates spherical micelles coexisting with longer, more flexible worm-like micelles (Supplemental Figure 3 in Supplemental Information).^34^ Given that enhanced extravasation and endocytosis are expected for sub-100 nm, nonspherical particles^39, 47^, CLNPs comprised of more than 50 mol% MVL5, with particle populations containing anisotropic micelles hold promise as hydrophobic drug delivery vectors. However, the viability of CLNPs containing MVL5 for PTX delivery applications has not been explored.

**Figure 1.**
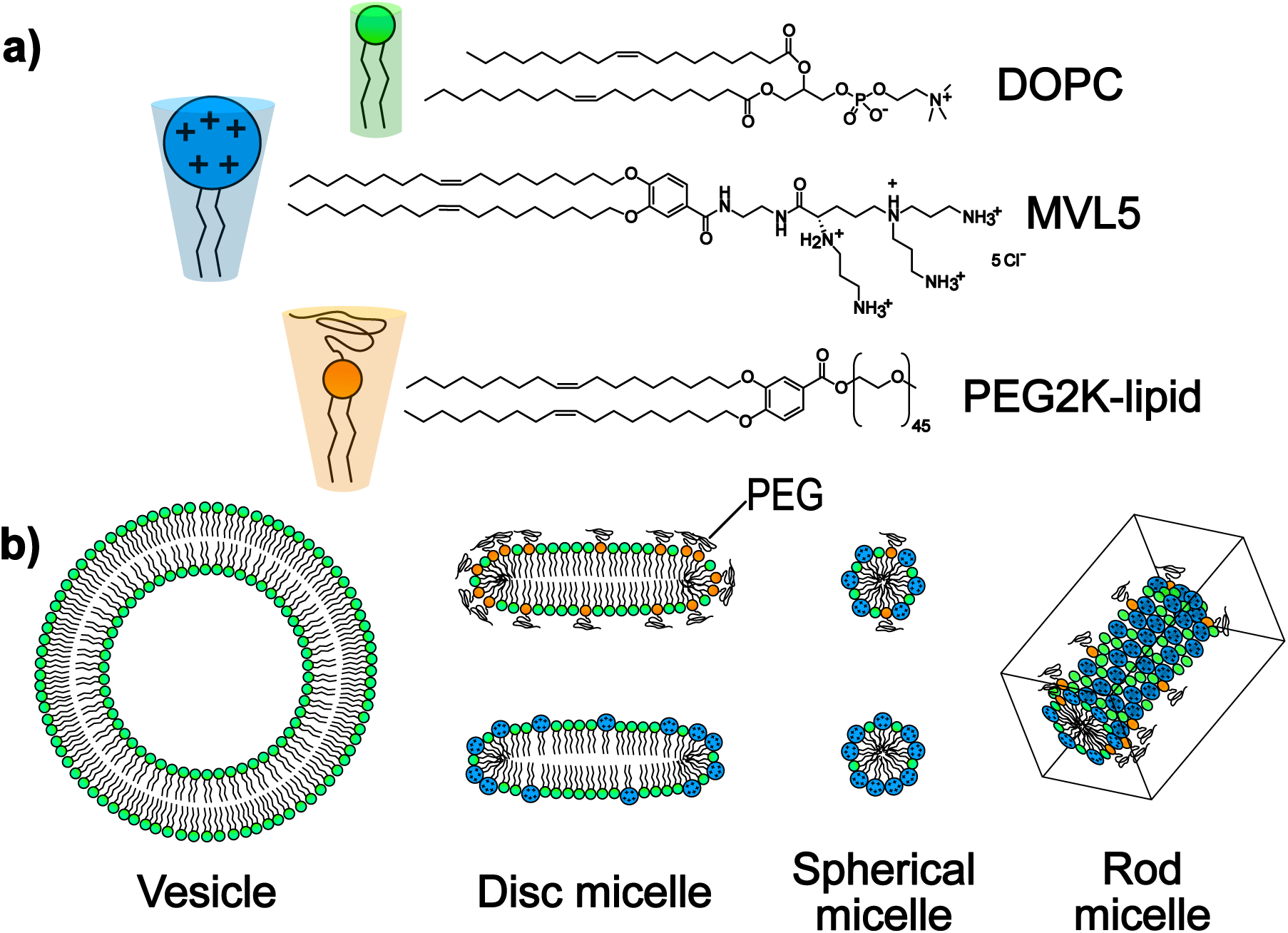
The structure and curvature of lipids investigated in this study and their relationship with nanoparticle morphology. The molecular structure of the neutral curvature lipid DOPC and positive curvature lipids MVL5 and 2kDa polyethylene(glycol) conjugated lipid (PEG2K-lipid) and representative images of the effective curvature of these lipids are shown in (a). Nanoparticle morphologies expected for formulations consisting of DOPC, DOPC and PEG2K-lipid, DOPC and MVL5, or DOPC, PEG2K-lipid, and MVL5 together are shown in (b).

In this study, we investigated the use of MVL5 CLNPs, with anisotropic and spherical fluid phase micellar morphologies, as PTX delivery vehicles. In particular, we characterized their PTX loading ability, PTX and drug-independent cytotoxicity, compatibility with PEGylation, and propensity for cellular uptake, with differential interference contrast (DIC) and fluorescence microscopy. The fluid phase (chain-melted) micelles studied here are expected to show significantly enhanced PTX solubility compared to solid phase (chain-ordered) anisotropic micellar structures of prior studies^14, 21, 38, 48–53^. To determine the PTX loading capacity of MVL5-containing CLNPs, we mapped out kinetic phase diagrams (KPDs) by using DIC microscopy to detect insoluble PTX crystallization in CLNPs containing 10 to greater than 82 mol% MVL5 with and without 10 mol% PEG-lipid and compared these to a DOTAP-containing CL formulation approximating the benchmark EndoTAG-1^TM^ formulation. We discovered an MVL5-dependent and positive curvature-dependent improvement in PTX solubility, where CLNPs with 50 mol% MVL5 and 50 mol% DOPC consisting of disc CLNPs (Supplemental Figure 2 in Supporting Information) solubilized 2-fold more PTX (from 3 to 6 mol%) than those with 50 mol% DOTAP and 50 mol% DOPC consisting primarily of vesicles (Supplemental Figure 1 in Supporting Information). Remarkably, KPDs at 75 mol% MVL5, where disc CLNPs transition to a coexisting mixture of spherical and short rod micelles (Supplemental Figure 2 in Supporting Information), exhibited improved solubility with PTX solubility extended to 8 mol% PTX for between 10 and 25 hours. PEGylation of CLNPs with micellar morphologies (50, 75, and 100−X mol% MVL5) primarily resulted in reduced time for PTX solubility at the larger PTX mol% range.

Cytotoxicity assays revealed that this improved PTX loading capacity translated into enhanced PTX-dependent cytotoxicity with reduced lipid toxicity for MVL5 compared to DOTAP CLNPs. Significant reduction in IC50 (the concentration of PTX at which cell viability was reduced to half the initial viability) was observed for MVL5 CLNPs compared to DOTAP CLNPs at 4 mol% (from 51 to 13 nM) and 3 mol% (from 17 nM to 11nM) PTX, which are at the solubility limit for DOTAP CLNPs but in the fully soluble regime for MVL5 CLNPs. The same assays revealed that MVL5 was much less toxic than DOTAP, which shows cytotoxicity at concentrations above 50 μM total lipid. PEGylation of MVL5 vehicles improved their cytotoxic efficacy to the same degree as DOTAP vehicles at the PTX solubility limit for each.

Particle analysis of fluorescent microscopy images of cancer cells treated with labeled MVL5 and DOTAP-containing CLNPs showed that PEGylation significantly improved cellular uptake and cell penetration depth, which was corroborated by fluorescent imaging of similarly treated live cells. For bare CLNPs, large fractions formed clumps bound to the cell surface. The stark contrast in cell uptake between bare and PEGylated CLNPs provides evidence of effective steric stabilization of the latter. This analysis also revealed a pattern of CLNP distribution that suggests the rate-limiting step for NP cell penetration and PTX delivery is transport of CLNP-containing endocytic vesicles through the actin mesh barrier near the cell surface and the rate at which PTX hops from NP to endosomal vesicle membrane and to microtubules.

## MATERIALS AND METHODS

### Materials

Lipid stock solutions of DOPC, DOTAP, MVL5, and 5 kDa PEG-DOPE were purchased from Avanti Polar Lipids as powder which was dissolved in chloroform to 10 mM. Unlabeled paclitaxel (PTX) was purchased from Fisher Scientific as a powder and dissolved at 10 mM in chloroform, OregonGreen 488 conjugated paclitaxel (OG-PTX) and TRITC-DHPE were also purchased from Fisher Scientific pre-dissolved in chloroform at 2 mM and 0.81 mM, respectively. Janelia Fluor 646® conjugated paclitaxel (Janelia-PTX) was purchased from Tocris Bioscience pre-dissolved in chloroform to 0.71 mM. CellTiter 96® AQueous-One Solution Cell Proliferation Assay was obtained from Promega. ProLong Gold Antifade Mountant with DAPI DNA Stain was purchased from Fisher Scientific.

### Cationic lipid nanoparticle (CLNP) preparation

Solutions of lipid and PTX were mixed in a chloroform:methanol (3:1, v/v) solvent in 1.5 mL glass vials at 1 mM total for all experiments. The solvent was evaporated under a nitrogen stream for 7 minutes, then the lipid mixture was further dried in a vacuum for 16 hours. The resultant film was resuspended in high resistivity (18 MΩ cm) water to 1 mM and taken directly for PTX solubility experiments, or sonicated then diluted

### Cell culture

The human prostate cancer cell line (PC3, ATC number: CRL-1435) and melanoma cell line (M21, subclone of UCLA-SO-M21 originally derived from the Reisfeld lab at Scripps Institute in La Jolla) were gifts from the Ruoslahti lab (Burnham Institute in La Jolla). Cells were cultured in Dulbecco’s modified Eagle’s medium (DMEM; Invitrogen) supplemented with fetal bovine serum (FBS; Corning) at 10% v/v and penicillin and streptomycin (Gibco) at 1% v/v. Cells were cultured in T75 flasks (Corning) at 37°C in a humidified incubator with 5% CO_2_ and split at a 1:5 ratio after reaching >80% confluence, around every 48 hours, during maintenance.

### Differential interference contrast (DIC) microscopy for PTX solubility assays

For PTX solubility experiments, 2 μL of rehydrated, PTX-loaded CLNPs stored at room temperature (around 22°C) were deposited into parafilm cutouts on glass slides, covered by a coverslip, and imaged at 20X magnification on an inverted Ti2-E microscope (Nikon) under DIC mode. Images were taken every 2 hours up to 12 hours, then every 12 hours up to 72 hours. Kinetic phase diagrams show the time at which crystals were present in samples of CLNP formulation at each mol% of PTX.

### Cell Viability Assays

96-well plates were seeded with 5,000 cells/well from a 50,000 cell/mL suspension in FBS supplemented culture media. Cells were allowed to adhere overnight. Sonicated CLNPs were diluted into DMEM to the desired concentration of PTX. Culture media was then replaced with 100 μL of the CLNP suspension and the CLNP treated cells were incubated for 24 hours. Then, the CLNP suspension was removed and replaced with culture media and cells were further incubated for 48 hours. At that point, cell viability was measured using CellTiter 96 Aqueous-One Solution Cell Proliferation Assay (Promega). The assay solution was diluted 1 to 3 in DMEM while the cell culture media in wells was replaced with 60 μL of DMEM, then 60 μL of the diluted assay solution was added rapidly to all wells via multichannel pipette. After around 1 hour incubation, 490 nm absorbance was measured using a plate reader (Tecan M220). Data points represent the average absorbance, shown as a percent of the absorbance obtained for untreated cells, for four identically treated wells at each PTX concentration for IC50 experiments on bare CLNPs and eight identically treated wells for each CLNP formulation tested for bare versus PEGylated CLNP experiments.

The IC50 for each PTX-loaded CLNP formulation was determined by finding the best fit line for the data in Python (curve_fit function) using y = A + (B-A)/(1+(x/C)^D^) as the fit equation. Here y is the cell viability at each PTX concentration, x is the concentration of PTX, A is the maximum cell viability, B is the minimum cell viability, C is the IC50 value for the curve, and D is the slope factor. To determine statistical significance between points on cell viability curves curves, a Student’s t-test was run using the average and standard error of measured cell viability for both formulations at each PTX concentration tested in excel. The same test was used to compare bare versus PEGylated CLNPs. Plotting was also done in excel.

### Fluorescence and DIC microscopy

For fixed cell imaging, M21 cells were seeded on poly-L-lysine coated glass cover slips in the wells of 6-well plates (Corning) by adding 2 mL of a 25,000 cell/mL suspension in culture media and allowing cells to adhere to cover slips overnight. The following day, culture media was aspirated, cells were washed once with PBS, and then 1 mL of sonicated CLNPs containing TRITC-DHPE and Janelia-PTX labels diluted in DMEM to 100 nM Janelia-PTX was added to the cells. Cells were incubated with the labeled particle solution for 6 hours then washed 3 times with PBS before being fixed and mounted using mounting media that included DAPI for nuclear staining. Images were taken on an inverted Ti2-E microscope (Nikon) at 90X magnification (60X lens with additional 1.5X magnification lens) in fluorescence and DIC mode.

For live cell imaging, M21 cells were seeded in glass-bottom 6-well plates (Corning) by adding 2 mL of a 50,000 cell/mL suspension in culture media and allowing cells to adhere overnight. Sonicated CLNPs containing TRITC-DHPE and OG-PTX labels suspended in DMEM at 400 nM OG-PTX were then added to the cells and incubated for 24 hours. Cells were washed 3 times with PBS before returning culture media and imaging in a chamber extension (OKO; H101- NIKON-TI-S-E, 6MW insert) held at 37°C and 5% CO_2_ on an inverted Ti2-E microscope (Nikon) at 20X magnification.

### Image Analysis

Quantitative analysis of the number of CLNP lipid puncta and puncta localization within cells was performed using ilastik and CellProfiler (version 4.2.7) image analysis software. Z-slices from Z-stack TIFF images that bisected the nucleus of all cells in the image frame were selected for analysis manually in ImageJ to ensure that puncta identified within the border of cells were inside the cell, rather than on the top or bottom surface. Z-slice TIFF images with all channels were imported into ilastik and a subset of images were used to train ilastik to identify cell borders across all images. Cell border traces identified by ilastik, and manually verified for de-clumping, as well was the original Z-slice TIFF images were imported into CellProfiler and used as input for a speckles analysis pipeline. This pipeline identified lipid puncta within cell borders then counted the number of puncta and their distances from the cell border from the input images shown in Supplemental Figures 4, 5, 6, and 7. Kruskal-Wallis and Dunn’s post-hoc test on puncta number and puncta-to-border distance data between each formulation tested was conducted in R using the tidyverse and rstatix packages. Plotting was also done in R.

## RESULTS AND DISCUSSION

### MVL5 Incorporation Improves the Solubility of Paclitaxel in Cationic Lipid Nanoparticle Membranes

Prior studies have shown that the solubility of PTX in CLNP membranes influences the cytotoxic efficacy of the PTX-loaded carrier^28^. Therefore, to determine the viability of CLNPs containing MVL5 as PTX delivery vehicles, we investigated the solubility of PTX in unsonicated CLNPs containing increasing MVL5 content both with 10 mol% PEG5k- DOPE (PEGylated) and without PEG5k-DOPE (bare). Owing to its hydrophobicity and propensity for self-association above a critical concentration, membrane insoluble PTX phase separates to form stable crystals which are visible under differential interference contrast (DIC) microscopy^28,48^. We used DIC microscopy to observe CLNP samples containing 2 to 8 mol% PTX at room temperature over time and identify when insoluble PTX crystals became visible in the sample. The kinetic phase diagrams (KPDs) in Figure 2 show how long after hydration PTX crystals were observed for every membrane PTX content in each CLNP formulation tested. For both bare and PEGylated CLNPs, increasing MVL5 content correlated with improved PTX membrane solubility. For bare CLNPs, the solubility limit increased from ≈3 mol% PTX for CLNPs containing 50 mol%

**Figure 2.**
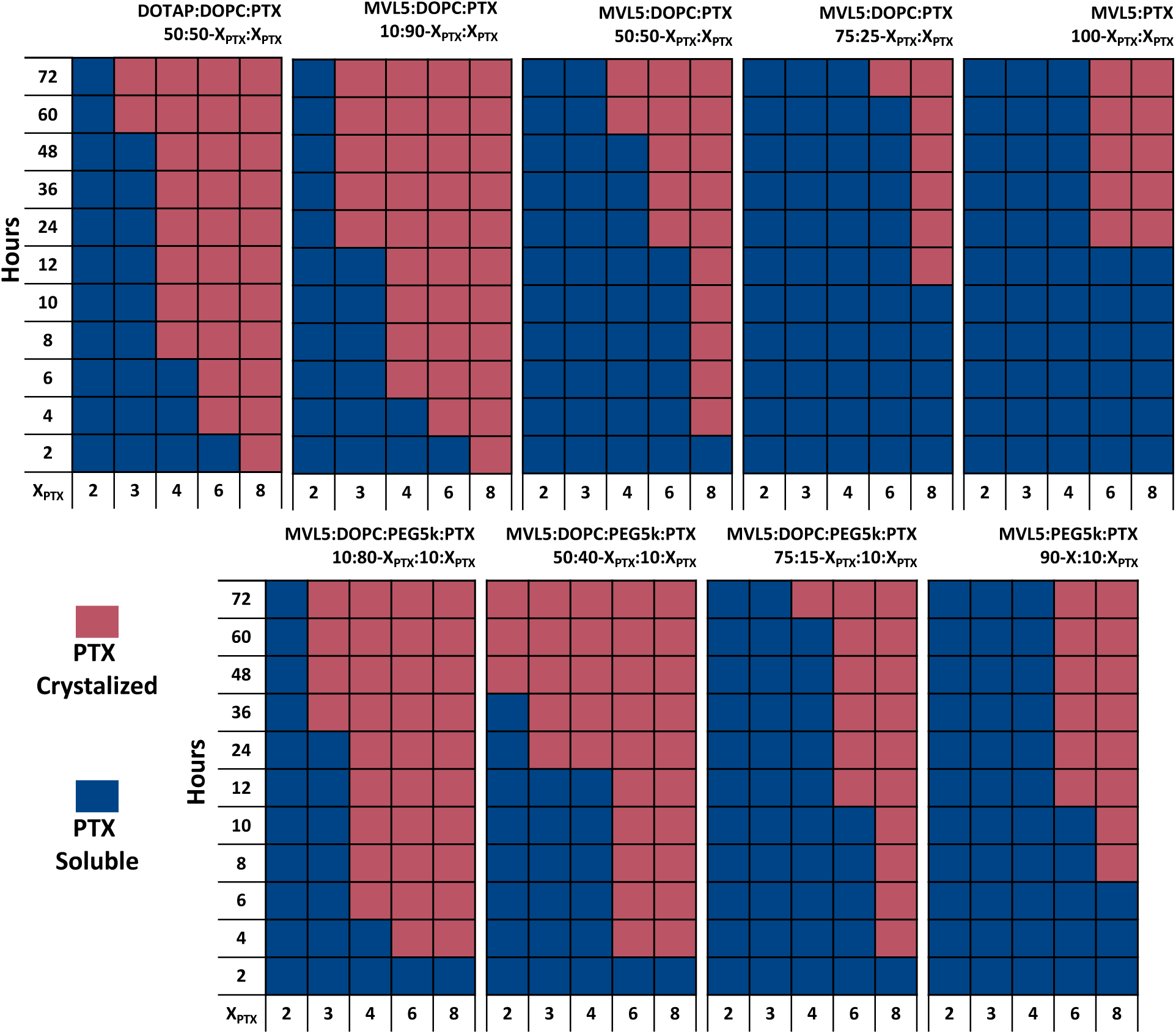
Solubility of paclitaxel (PTX) at varied concentrations in lipid membranes of unsonicated cationic liposomes (CLNPs) over time as a function of membrane MVL5 and PEG-lipid content. Diagrams indicate the time over which PTX remained solubilized in the membrane of CLNPs after lipid film hydration. Blue indicates that PTX remained solubilized in the liposome membrane while red indicates that PTX phase separated and precipitated out of the membrane, as determined by the presence of crystals in samples observed under DIC microscopy. CLNP formulations are shown above each diagram, with MVL5 content increasing from 10 to >92 mol % from left to right. Bare CLNP formulations are shown in the top row, with a CLNP formulation containing 50 mol% DOTAP shown on the far left for comparison. CLNP formulations containing 10 mol % PEG-lipid, which replaced DOPC or MVL5 when no DOPC was incorporated, are shown on the bottom row.

DOTAP or 10 mol% MVL5 to ≈6 mol% for CLNPs containing 75 mol% MVL5 or entirely MVL5 and PTX. Significantly, even at 8 mol% PTX, CLNPs with 75 mol% MVL5 or 92 mol% solubilized PTX up to 10 to 12 hours. The same trend was observed for PEGylated CLNPs, whose solubility limit also increased from ≈3 mol% PTX to ≈6 mol% PTX as MVL5 content increased from 10 to >80 mol%. However, the lifetime of solubility decreased, in particular, for 50 mol% MVL5 CLNPs at 6 mol% PTX.

Figure 3 shows select DIC images which display examples of PTX crystals among PTX-loaded CLNPs (for bare and PEGylated CLNPs) in insoluble PTX regimes and PTX-loaded CLNPs alone in soluble PTX regimes, and which were used to generate kinetic phase diagrams. The size and thickness of PTX crystals found in CLNP samples with high (75 to >90 mol%) MVL5 content were reduced due to the higher number of distinct PTX nucleation sites present in samples with higher quantities of smaller particles, similar to PTX crystals observed in PEGylated CLNPs here and in prior studies.^35^ This corresponds with the transition of CLNPs from spherical vesicles to micellar structures, especially spherical and worm micelles found at 75 mol% MVL5 and above for bare CLNPs (Supplemental Figure 2 in Supporting Information) and at 50 mol% MVL5 and above for PEGylated CLNPs in Cryo-TEM (Supplemental Figure 3 in Supporting Information).^34^ This transition can also be seen in the disappearance of visible vesicles under DIC microscopy seen in images of samples at or above those MVL5 contents (Figure 3, right panels).

**Figure 3.**
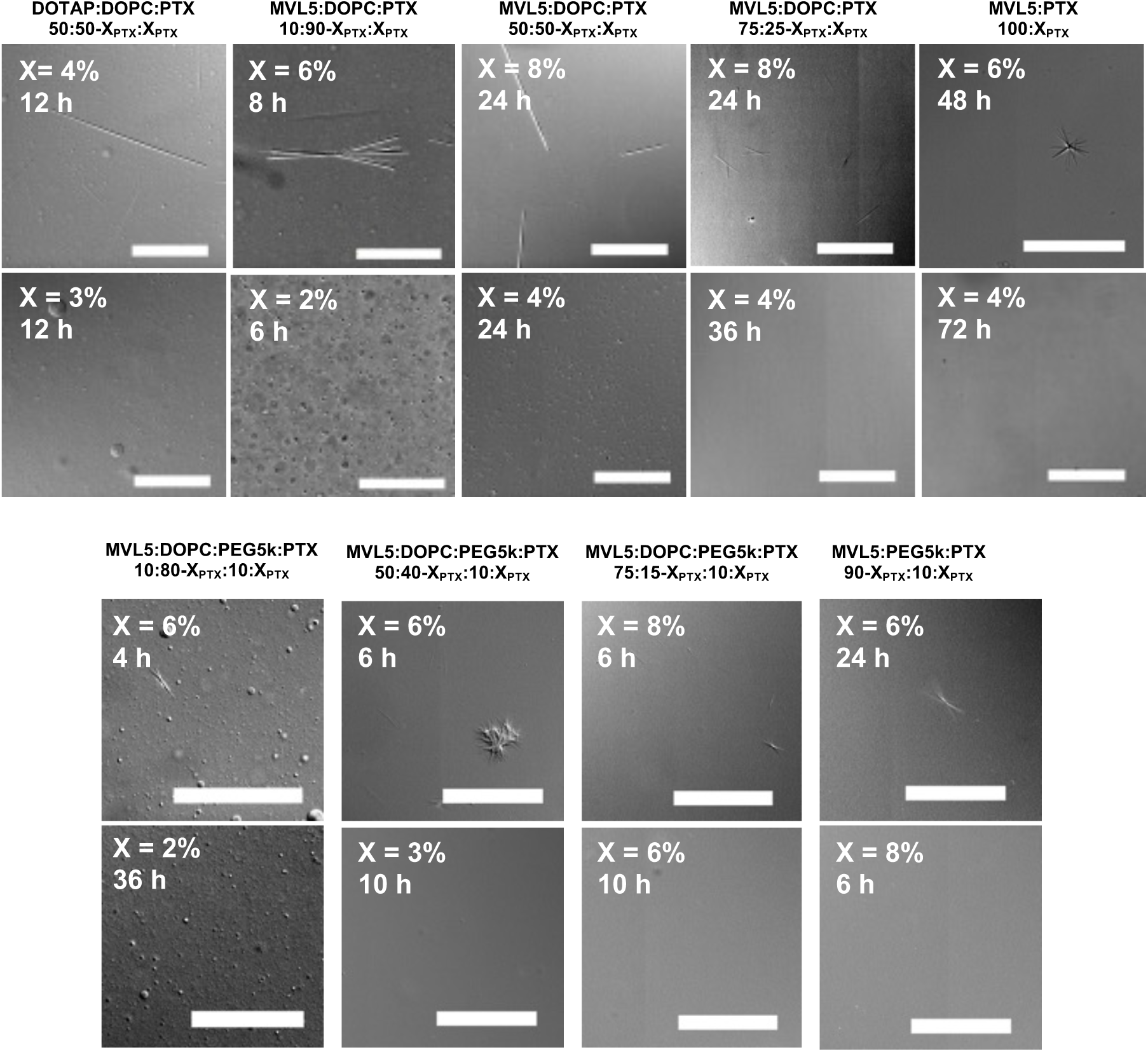
Differential interference contrast (DIC) microscopy images of cationic liposomes (CLNPs) and paclitaxel (PTX) crystals. Samples in the top row are unsonicated liposomes made of MVL5, DOPC, and PTX except for the leftmost panels, where they are made of DOTAP, DOPC, and PTX, at the molar ratios specified in the figure. Samples in the bottom row are unsonicated liposomes made of MVL5, DOPC, PEG5k-DOPE, and PTX at the molar ratios specified in the figure. The formulations used in the top and bottom rows of this figure correspond to the formulations used in the top and bottom rows, respectively, of Figure 1. The time since hydration of the lipid film is indicated on each image. Scale bars are 100 µm.

### Paclitaxel Solubility Correlates with the Average Spontaneous Curvature of Membranes

Previous work has shown that lipids with negative spontaneous curvature (i.e. lipids with inverse conical shapes that form inverse hexagonal, or HII phase, membranes) have reduced PTX solubility compared to those with neutral spontaneous curvature (i.e. lipids with cylindrical shapes that form lamellar, or L⍺ phase, membranes)^54^. To explore the influence of curvature on PTX solubility, we generated kinetic phase diagrams for bare and PEGylated CLNP formulations containing the negative spontaneous curvature lipid DOPE (i.e. with headgroup smaller than the double tails), and either the neutral or positive spontaneous curvature cationic lipids DOTAP or MVL5, respectively. The spontaneous curvature of the CLNP membranes is expected to transition from negative for formulations containing >72 mol% DOPE and 20 mol% DOTAP towards being less negative and crossing zero curvature in formulations containing MVL5 and DOPE as their MVL5 content increases from 10 to 50 mol% MVL5.

Figure 4 shows the kinetic phase diagrams generated from DIC imaging of PTX crystal formation over time for these formulations. This transition from negative to positive average spontaneous curvature coincided with improved PTX solubility, with the solubility limit increasing from 1 to ≈3 mol% PTX as the cationic lipid content shifts from 20 mol% DOTAP to 50 mol% MVL5 for both bare and PEGylated CLNPs. The reduced particle size in DIC microscopy images of CLNP formulations containing 50 mol% MVL5 shown in Figure 5 provides evidence that the expected curvature shift from negative to positive indeed occurred. For bare CLNPs, but not for PEGylated CLNPs, the size of observed PTX crystals also reduced with the transition from negative to positive curvature.

**Figure 4.**
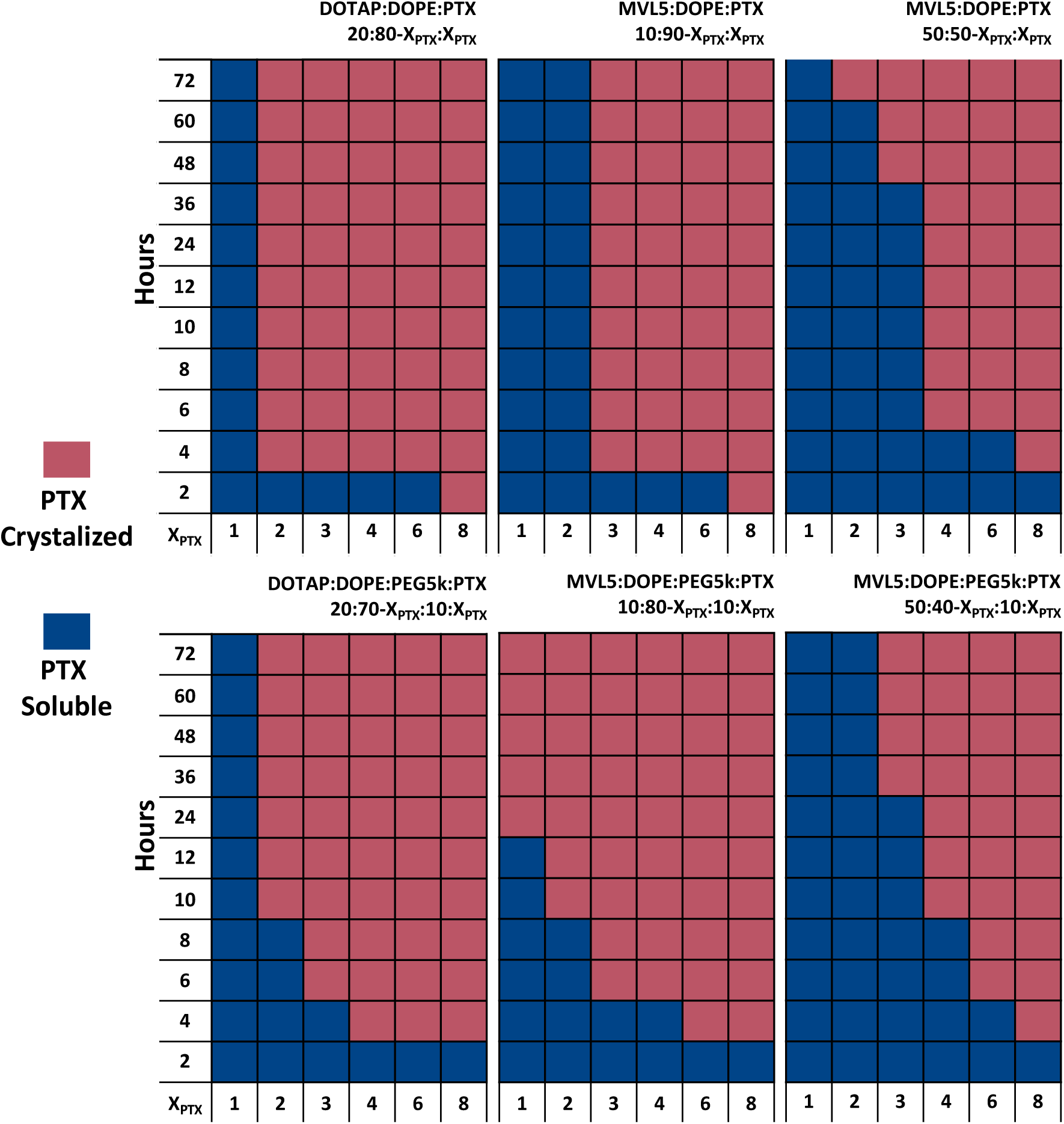
Solubility of paclitaxel (PTX) in lipid membranes of unsonicated, cationic liposomes (CLNPs) over time as a function of membrane curvature. Diagrams indicate the time over which PTX remained solubilized in the membrane of CLNPs after lipid film hydration. Blue indicates that PTX remained solubilized in the liposome membrane while red indicates that PTX phase separated and precipitated out of the membrane, as determined by the presence of crystals in samples observed under DIC microscopy. CLNP formulations are shown above each diagram, with the expected curvature of the CLNP membranes progressing from highly inverted (containing 80 mol % DOPE, top left diagram) to a more neutral curvature (containing 50 mol % MVL5 and >42 mol % DOPE) from left to right. Bare CLNP formulations are shown on the top row while CLNP formulations containing 10 mol % PEG-lipid, which replaced DOPE lipid, are shown in the bottom row.

**Figure 5.**
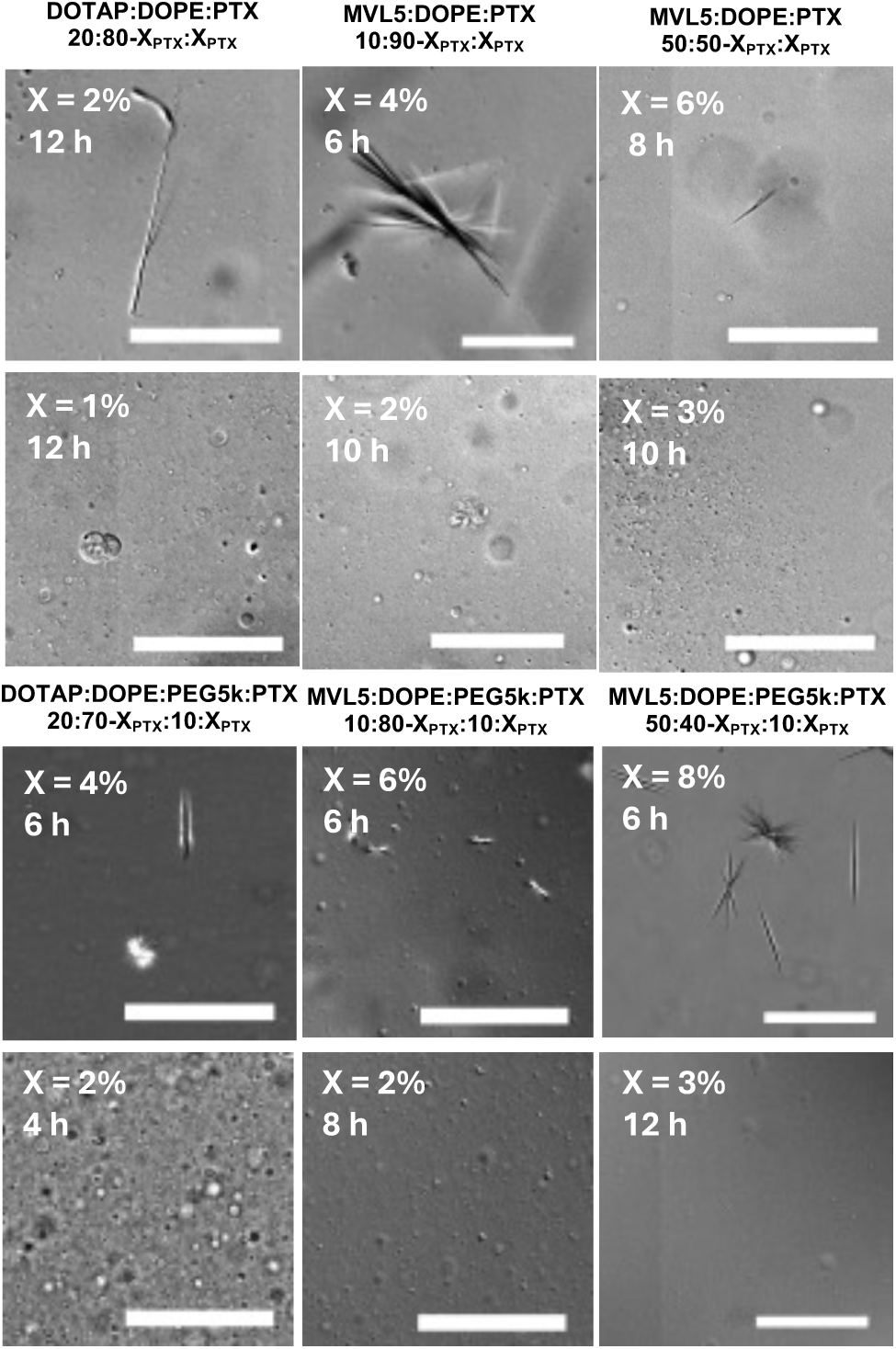
DIC images of cationic liposomes with solubilized or insoluble, crystalized PTX. Samples in the top row are unsonicated liposomes comprised of DOTAP, DOPE, and PTX or MVL5, DOPE, and PTX at the molar ratios specified in the figure. The samples in the bottom row are unsonicated liposomes comprised of DOTAP, DOPE, PEG5k-DOPE, and PTX or MVL5, DOPE, PEG5k-DOPE, and PTX at the molar ratios specified in the figure. The formulations used in the top and bottom rows of this figure correspond to the formulations used in the top and bottom rows, respectively, of Figure 2. The time after lipid film hydration is indicated on each image. Scale bars are 100 µm.

### Improved Paclitaxel Solubility of Cationic Lipid Nanoparticles Containing MVL5 Leads to Higher Cytotoxic Efficacy *In Vitro*

We investigated the ability of sonicated, bare CLNPs containing 50 mol% DOTAP or MVL5, (50-X_PTX_) mol% DOPC, and 2, 3, or 4 mol% PTX membrane content to deliver PTX to human prostate cancer (PC3) cells using cell viability assays.

The kinetic phase diagram data show that 2 to 4 mol% PTX crosses from below to above the PTX solubility limit of DOTAP-CLNPs, while remaining well below (for 50 mol% and greater MVL5 content) or at the PTX solubility limit of MVL5-CLNPs (for 10 mol% MVL5 content). Therefore, these assays elucidated two points: first, whether the improved PTX solubility observed in MVL5- CLNPs translates into higher cytotoxic efficacy; and second, for the assay run at 2 mol% PTX, where both DOTAP- and MVL5-CLNPs solubilize PTX, whether MVL5 confers higher cytotoxic efficacy due to factors other than PTX solubility, such as altered particle shape and size.

Figure 6 shows the results of these cell viability assays and the IC50 of each CLNP formulation derived from those results, as well as the cell viability of PC3 cells after treatment with DOTAP- and MVL5-CLNPs without PTX loading. At 4 mol% PTX membrane content, the cytotoxic efficacy of MVL5-CLNPs was higher than DOTAP-CLNPs, shown by the more rapid loss of cell viability at lower PTX concentrations (Figure 6a) and the ≈3.9-fold lower IC50 (Figure 6e). At 4 mol% PTX content, PTX in DOTAP-CLNPs phase separates into insoluble crystals within 24 hours, the time cells were exposed to particles in this assay, versus 60 hours for MVL5-containing particles. At 3 mol% PTX, where DOTAP-CLNPs are at their solubility limit while MVL5-CLNPs are below their limit, the cytotoxic efficacy of MVL5-CLNPs remains higher but the difference in efficacy is reduced, with the IC50 of MVL5-CLNPs only ≈1.5-fold lower (Figure 6b,e). The difference in cytotoxicity collapsed at 2 mol% PTX, where both DOTAP- and MVL5-CLNPs are below their solubility limit, with IC50 equal to 12.2 and 10.4 nM PTX for MVL5- and DOTAP-CLNPs, respectively (Figure 6c,e). These results demonstrate that improved PTX solubility in MVL5-CLNPs enhances their cytotoxic efficacy compared to DOTAP-CLNPs at PTX contents greater than 3 mol%, enabling effective delivery of higher PTX doses at equal lipid concentrations *in vitro* and *in vivo*. Further, the <30 nm size and disc micelle structure of MVL5-CLNPs may enhance extravasation and tumor accumulation *in vivo*, which was observed in prior studies of PEGylated DOTAP-CLNPs with similar structures^40^.

**Figure 6.**
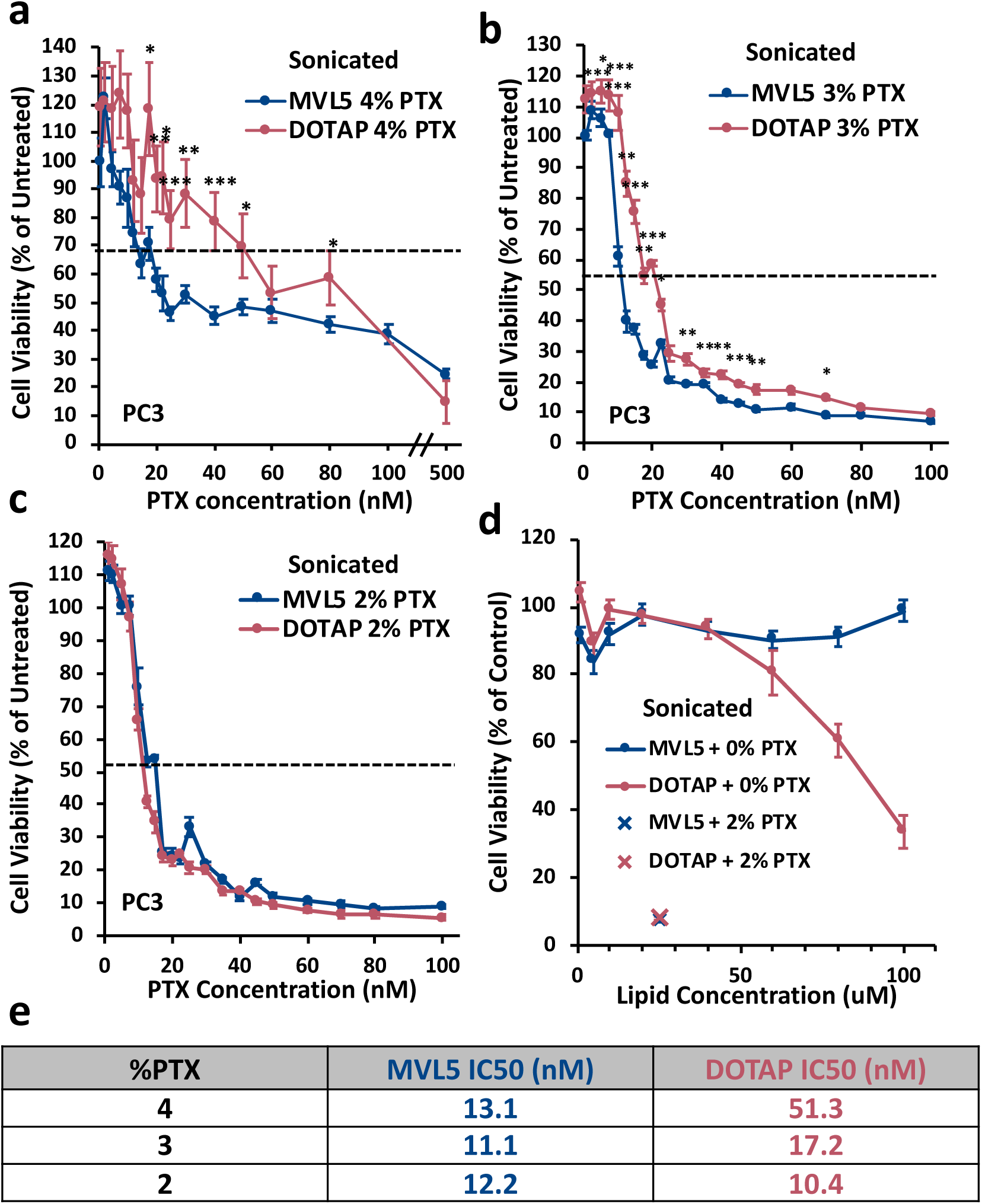
Viability of PC3 cells, compared to untreated control cells, as a function of PTX concentration for cells treated with PTX-loaded, sonicated, bare CLNP formulations. The molar compositions of these CLNPs were MVL5/DOPC/PTX equal to 50/50−X_PTX_/X_PTX_ mol % (blue line with circles) and DOTAP/DOPC/PTX equal to 50/50−X_PTX_/X_PTX_ mol % (red line with circles) where formulation PTX content was 4 (**a**), 3 (**b**), 2 (**c**), and 0 or 2 (**d**) mol %. Dashed black lines in a, b, and c indicate the cell viability which is halfway from the maximum and minimum viability measured for each formulation pair, which was taken for IC_50_ determination, which is summarized in (**e**). The blue vertical and red diagonal crosses in figure (**d**) show the viability of cells treated with MVL5 and DOTAP CLNPs, respectively, loaded with 2 mol % PTX (final PTX concentration applied to cells = 500nM), measured independently from the experiment shown in (**c**). Error bars indicated standard error across 4 replicates. Statistical significance was determined by Welch’s T-test for each PTX concentration and is indicated by asterisks: (*) for P < 0.05, (**) for P < 0.01, and (***) for P < 0.001.

Figure 6d shows the results of an experiment which investigated the intrinsic toxicity of MVL5 compared to DOTAP using a cell viability assay. PC3 cells were treated with MVL5- and DOTAP-CLNPs which were not loaded with PTX at total lipid concentrations of 0 to 100 µM or treated with the same CLNPs loaded with 2 mol% PTX at 25 µM total lipid (corresponding to 500 nM PTX). MVL5-CLNPs without PTX loading showed no cytotoxicity, as cell viability remained within 10 % of untreated control cells across all lipid concentrations tested except one outlier at 5 µM total lipid. In contrast, cell viability for DOTAP-CLNPs without PTX fell to ≈80 % of control cells at 60 µM total lipid and continued to decline to ≈34 % of control cells by 100 µM total lipid, demonstrating that the innate toxicity of DOTAP was detected in this assay. However, this toxicity occurs at total lipid concentrations well beyond those required to induce cytotoxicity when loaded with 2 mol% PTX, as PTX-loaded DOTAP- and MVL5-CLNPs added at only 25 µM total lipid reduced cell viability to ≈8 and ≈7 % of untreated control cells, respectively. Nonetheless, the lack of innate cytotoxicity for MVL5 at any lipid concentration tested means it would have reduced off target effects compared to the established cationic lipid, DOTAP, when administered at therapeutically relevant total lipid concentrations of 50 mg/mL (≈67mM), which are required to achieve sufficiently high PTX dosage for chemotherapy^55, 56^.

### PEGylation of Cationic Lipid Nanoparticles Containing MVL5 Further Improves Cytotoxic Efficacy *In Vitro*

The inclusion of PEG5k-DOPE above 10 mol%, where the PEG polymer is forced into a linear conformation and is thus termed the brush regime, drives the alteration of DOTAP-containing PTX-loaded CLNP shape and size into <30 nm disc micelles and enhances their cellular uptake and cytotoxic efficacy^35^. Since bare CLNPs containing 50 mol% MVL5 or greater form entirely disc micelles, we investigated whether the cytotoxic efficacy of these CLNPs was saturated at that particle structure profile, or if PEGylation would further improve cytotoxic efficacy through additional shape changes or other alterations, such as steric stabilization. Figure 7 presents cell viability assay results for PC3 cells treated with sonicated MVL5- and DOTAP-CLNPs loaded with 4 and 2 mol% PTX, respectively, which is just below the solubility limit of each formulation, with and without 10 mol% PEG5k-DOPE, at a fixed PTX concentration of 20 nM. PEGylation similarly enhanced the cytotoxic efficacy of both MVL5- and DOTAP-CLNPs, reducing cell viability by 41.8% and 36.6%, respectively, compared to cells treated with bare formulations. This finding suggests that PEGylation at or above 10 mol% PEG5k-lipid enhances cytotoxicity through mechanisms other than, or in addition to, driving the formation of disc micelles. Since Cryo-TEM data of CLNPs at 50 mol% MVL5 show that PEGylation transforms particle structure from disc micelles to spherical and elongated worm micelles (Supplemental Figure 2 and Supplemental Figure 3 in Supporting Information),^34^ it is possible that this additional change in CLNP morphology confers improved cellular uptake, and therefore cytotoxic efficacy, compared to disc micelles. PEGylation in the brush regime also significantly bolsters the steric stabilization of MVL5-CLNPs and prevents aggregation following accumulation of particles at the cell surface due to sulfated glycosaminoglycan binding, which would limit cellular uptake.

**Figure 7.**
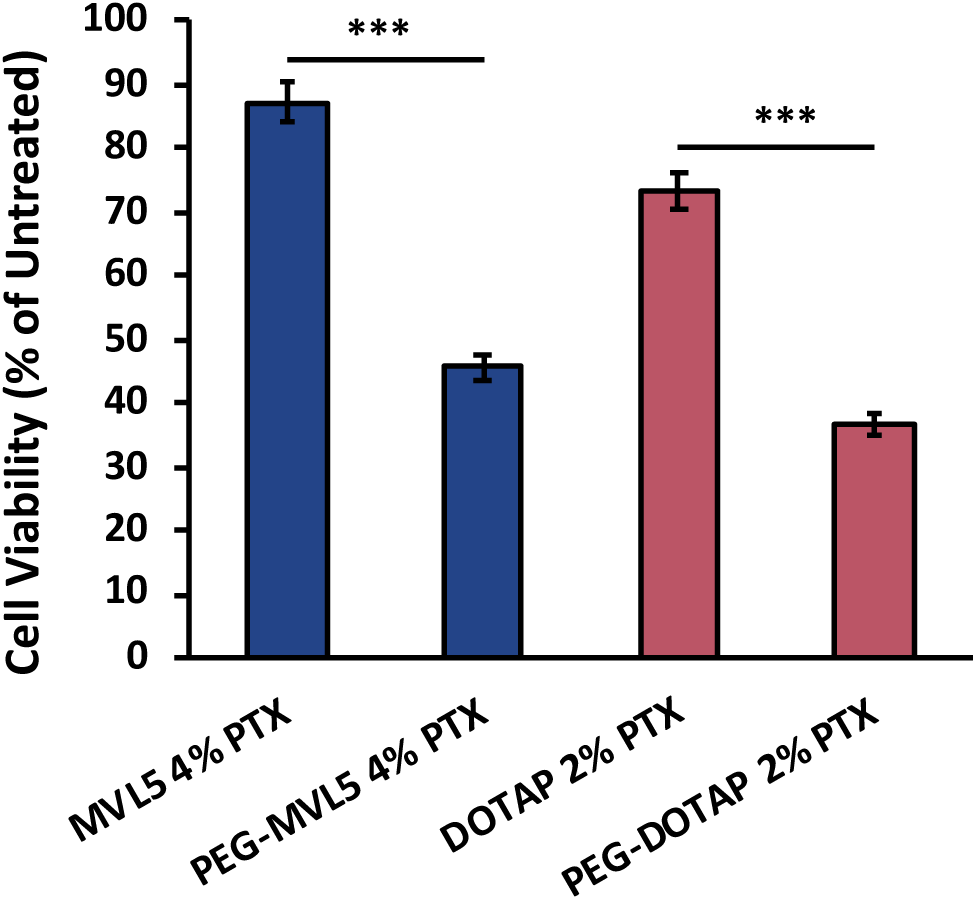
Viability of PC3 cells treated with bare and PEGylated CLNPs loaded with PTX at a fixed PTX concentration of 20 nM compared to untreated control cells. CLNP molar composition was 0/10 mol % PEG-lipid, 50 mol % MVL5, 46−X_PEG_ mol % DOPC, and 4 mol% PTX (blue bars) and 0/10 mol % PEG-lipid, 50 mol % DOTAP, 48−X_PEG_ mol % DOPC, and 2 mol% PTX (red bars). Error bars represent the standard error for viability measurements across 8 replicates. Statistical significance determined by Welch’s T-test between bare and PEGylated CLNPs is indicated by asterisks: (*) for P < 0.05, (**) for P < 0.01, and (***) for P < 0.001.

### PEGylation of Cationic Lipid Nanoparticles Containing MVL5 Promotes Enhanced Cellular Uptake

To determine whether the enhanced cytotoxic efficacy of PEGylated MVL5- CLNPs was related to improved cellular uptake, we used fluorescence microscopy of fixed melanoma cancer (M21) cells treated with sonicated, 50 mol% MVL5 CLNPs containing fluorescent lipid (TRITC-DHPE) and PTX (Janelia-PTX) labels with and without 10 mol% PEG5k-DOPE. Select X-Y plane images at chosen Z intersecting the bulk of the cell body including the nucleus (shown in blue), as well as Z-Y and Z-X plane views, from fluorescence microscopy images of representative cells treated with bare and PEGylated MVL5-CLNPs are shown in Figure 8. Lipid (shown in green) in bare MVL5-CLNPs (Figure 8a) accumulated near the border of cells, often in large aggregates, while PTX (shown in red) diffused into the body of the cell. In contrast, lipid in PEGylated MVL5-CLNPs (Figure 8c) did not show aggregation, and instead remained as separate puncta (small, discrete spots in green). Lipid from PEGylated MVL5-CLNPs also showed some accumulation near the cell surface but showed significant uptake into the cell body, with puncta distributed throughout the cross-sectional area of the cell body, from the cell surface to the nucleus. These MVL5-CLNPs were compared to controls consisting of identically labeled, sonicated CLNPs at 50 mol% DOTAP both with (Figure 8d) and without (Figure 8b) 10 mol% PEG5k-DOPE. DOTAP-CLNPs showed similar patterns of uptake where bare DOTAP-CLNPs accumulated in large aggregates at the cell surface while PEGylated DOTAP-CLNPs had significantly increased uptake into the cell body as separate puncta. The enhanced uptake with PEGylation of CLNPs containing DOTAP and DOPC was also reported previously.^35^

**Figure 8.**
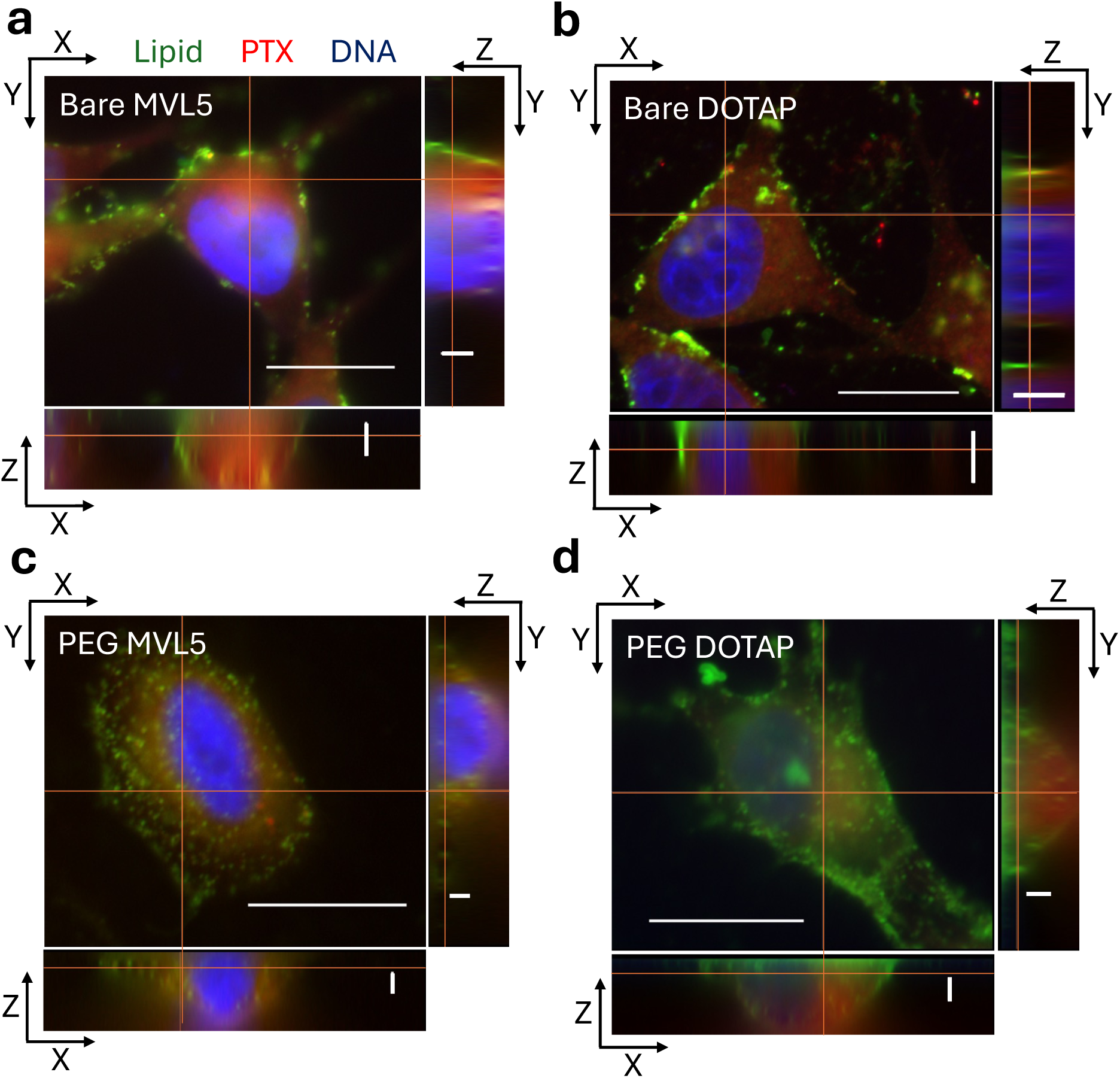
X-Y plane views at selected Z from a Z-stack image series and Z-Y and Z-X plane views from fluorescent microscopy images of fixed M21 cells treated with bare and PEGylated CLNPs containing fluorescently labeled lipid and PTX-fluorophore conjugate. M21 cells were treated with CLNPs of molar composition DOTAP/DOPC/PEG-lipid/TRITC-DHPE/Janelia-PTX equal to 50/46.5−X_PEG_/X_PEG_/0.5/3 mol% with either 0 (**a**) or 10 (**c**) mol% PEG-lipid and CLNPs with composition MVL5/DOPC/PEG-lipid/TRITC-DHPE/Janelia-PTX equal to 50/46.5−X_PEG_/X_PEG_/0.5/3 mol% with 0 (**b**) or 10 (**d**) mol% PEG-lipid. CLNP lipid (TRITC-DHPE) signal is shown in green, PTX (Janelia-PTX) signal is shown in red, DAPI was used to stain cell nuclei after fixation and is shown in blue. The main image at the top left panel of each image set shows an X-Y section at a selected Z from a Z-stack image series which intersects the body of the cell. The right projection shows the Y-Z side view of the slice along the vertical orange line in the main image. The bottom projection shows the X-Z side view of the slice along the horizontal orange line in the main image. Scale bars on the main image and projections are 20 mm and 2.5mm, respectively.

### Image Analysis of Lipid Puncta in Cells

To quantify CLNP cell uptake and penetration, we used an automated image analysis program that identifies the cell border and lipid puncta, counts the number of lipid puncta within the cell border, and measures their distance from the nearest point on the cell border (see materials and methods). Examples of images used for analysis, which in this case are lower magnification images of the same X-Y planes at selected Z shown in Figure 8, and the results of the puncta count and puncta to border distance analysis, are shown in Figure 9. Figure 9a shows the channels corresponding to fluorescently labeled CLNP-lipid, PTX, and DNA separated by column in green, red, and blue, respectively, along with a DIC and merged fluorescence image for each formulation. Qualitatively, these images show the same trend across larger populations of cells as seen in the representative cells from Figure 8, with bare DOTAP- and MVL5-CLNPs accumulated on the surface of cells and sparse lipid accumulation inside the cell body compared to PEGylated CLNPs (Figure 9a top and third row versus second and fourth row). Quantification confirmed this difference in cellular uptake of CLNPs, as the average number of identified CLNP-lipid puncta within the cell border of cells treated with PEGylated DOTAP- and MVL5-CLNPs was 2.6 and 3.4-fold higher, respectively, than the those treated with bare CLNPs containing the same lipid (Figure 9b). Taken together, these data show that inclusion of PEG-lipid at molar contents corresponding to the brush regime, where PEG-coated CLNPs are sterically stabilized, enhance the cellular uptake of both DOTAP- and MVL5-CLNPs to a similar degree.

**Figure 9.**
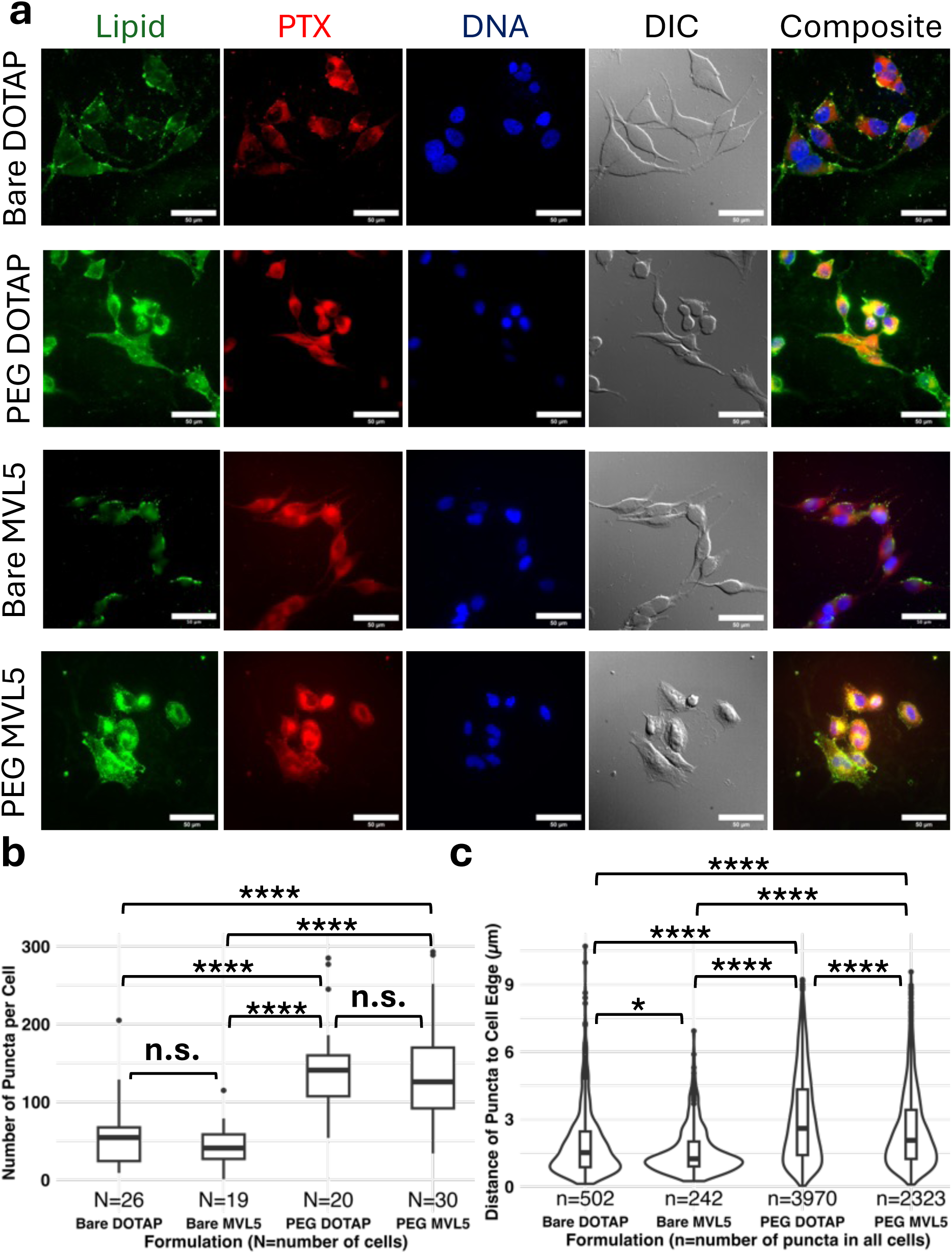
Selected z-slices from fluorescent microscopy image stacks of fixed M21 cells treated with bare and PEGylated CLNPs containing fluorescently labeled lipid and PTX-fluorophore conjugate. Images of M21 cells treated with CLNPs of molar composition MVL5/DOPC/PEG-lipid/TRITC-DHPE/Janelia-PTX equal to 50/46.5−X_PEG_/X_PEG_/0.5/3 mol % and DOTAP/DOPC/PEG-lipid/TRITC- DHPEE/Janelia-PTX equal to 50/46.5−X_PEG_/X_PEG_/0.5/3 mol % with either 0 or 10 mol % PEG-lipid are shown in (**a**). CLNP lipid (TRITC-DHPE) signal is shown in green, PTX (Janelia-PTX) signal is shown in red, DAPI was used to stain cell nuclei after fixation and is shown in blue. Scale bars are 50 μm.. The number of lipid puncta observed within the border of the cells treated with each formulation at this z-slice is quantified in (**b**) using the Cell Profiler particle (speckles) analysis. The distance of the identified lipid puncta from the cell border for cells treated with each formulation is quantified in (**c**). Statistical significance was determined using Dunn’s test in R and shown by asterisks (* for P<0.05, ** for P<0.01, *** for P<0.001, **** for P<0.0001). In (**b**), a box plot is shown where the ranked puncta per cell data for each formulation is binned into quartiles, the range of which are shown as vertical lines for the first and fourth quartiles, and the second and third quartiles are shown as a box, with the median of data shown as a horizontal line. Outliers were determined as points greater than 1.5 times the range of the second and third quartile away from the start of the fourth quartile and are shown as individual dots. In (**c**) a violin plot of the ranked distance of puncta to cell edge data, where a gaussian curve estimating the relative density of data points at each distance value along the y-axis is plotted, is overlayed on a box plot of the same data set.

Since populations of PEGylated DOTAP and bare, 50 mol% MVL5 CLNPs are either coexisting mixtures of vesicles and disc micelles (for DOTAP) or entirely or entirely disc micelles (for MVL5), respectively, the finding that only PEGylated DOTAP-CLNPs display enhanced uptake shows that steric stabilization is necessary to facilitate CLNP uptake regardless of size and morphology (smaller vesicles and micelles). The mechanism by which PEG in the brush layer enhances cellular uptake^35, 40^ is therefore likely through mechanisms beyond disc micelle formation, such as PEG-mediated steric stabilization, which limits aggregation of PEGylated CLNPs compared to those with PEG in the mushroom regime as well as bare CLNPs. The structural transition of 50 mol% MVL5 CLNPs from disc micelles to spherical and worm micelles through PEGylation was another possible driver of enhanced cellular uptake. However, if that was a major contributor to enhanced uptake, then the uptake of PEGylated MVL5-CLNPs should benefit from both structural and non-structural alterations, such as steric stabilization, and show improved uptake compared to PEGylated DOTAP-CLNPs, which was not observed.

Lipid puncta to cell-border distance analysis revealed a pattern of puncta distribution where the largest population of puncta was found near the cell surface, at around ≈1.6 µm from the cell border, but a long tail of the population reached distances of ≈6.9 to ≈10.7 µm in the cytoplasm from the cell border, 4.3 to 6.6-fold farther than the bulk of the population, in cells treated with all CLNPs tested (Figure 9c, all formulations). Despite this similarity in the pattern of puncta distribution, puncta populations for PEGylated CLNPs had significantly higher proportions of puncta at higher distances from the cell border compared to bare CLNPs. This is evident in the wider extended tails above 3 µm from the cell edge in population density violin plots of puncta in PEGylated CLNP-treated compared to bare CLNP-treated cells (Figure 9c, right two formulations versus left two formulations).

The distribution pattern and higher population of puncta beyond 3 µm from the cell edge for PEGylated CLNP-treated cells supports a model of CLNP uptake where the ≈100 to ≈200 nm mesh size of the actin network near the cell surface requires significant actin remodeling for endocytic vesicles containing CLNPs larger than 200 nm to reach microtubules.^57, 58^ However, once those endosomal vesicles reach a microtubule, they are rapidly brought far into the cell body via active transport. Figure 10 depicts this model of CLNP uptake and confinement in endosomal vesicles. The aggregation of bare CLNPs near the surface of cells likely means that they are endocytosed in vesicles which are larger than 200 nm. In contrast, the steric stabilization of PEGylated CLNPs prevents their aggregation, maintains smaller particle diameters, and facilitates packaging into vesicles under 200 nm. While the actin network mesh size restricts diffusion of endocytic vesicles carrying CLNPs, leading to the accumulation of both bare and PEGylated CLNP-carrying vesicles near the cell surface, the proportion of vesicles with PEGylated CLNPs that reach microtubules would be greater in this model. Active transport of those, and the few bare CLNP-containing vesicles which reach microtubules, then rapidly transports the CLNPs deep into the cell body (Figure 10a). CLNPs contained within endosomal vesicles may also alter the effective charge of endocytic vesicle membranes through attractive electrostatic interactions with the anionic endosomal vesicle membrane and cationic lipid hopping from CLNP to the endosomal vesicle membrane. The presence of cationic lipid domains on endosomal membrane may facilitate the adhesion of endosomal vesicles to microtubules (Figure 10b), a phenomenon which is observed for cationic liposomes and microtubules.^59^ In our proposed model, since the pharmacological target of PTX is microtubules, and the hydrophobic PTX molecule can hop from the CLNP membrane inside an endosomal vesicle to the endosomal membrane and eventually the β-subunit of microtubules once they are in proximity, crossing the actin network and PTX hopping from the endosomal vesicle to microtubules are likely the rate limiting steps in effective PTX delivery (this would apply for the case where the endosome is partly adhered to the microtubule or with the endosomal membrane tethered to the microtubule via a motor protein). This explains the marked improvement in cytotoxic efficacy seen with PEGylated CLNPs compared to bare CLNPs seen here.

**Figure 10.**
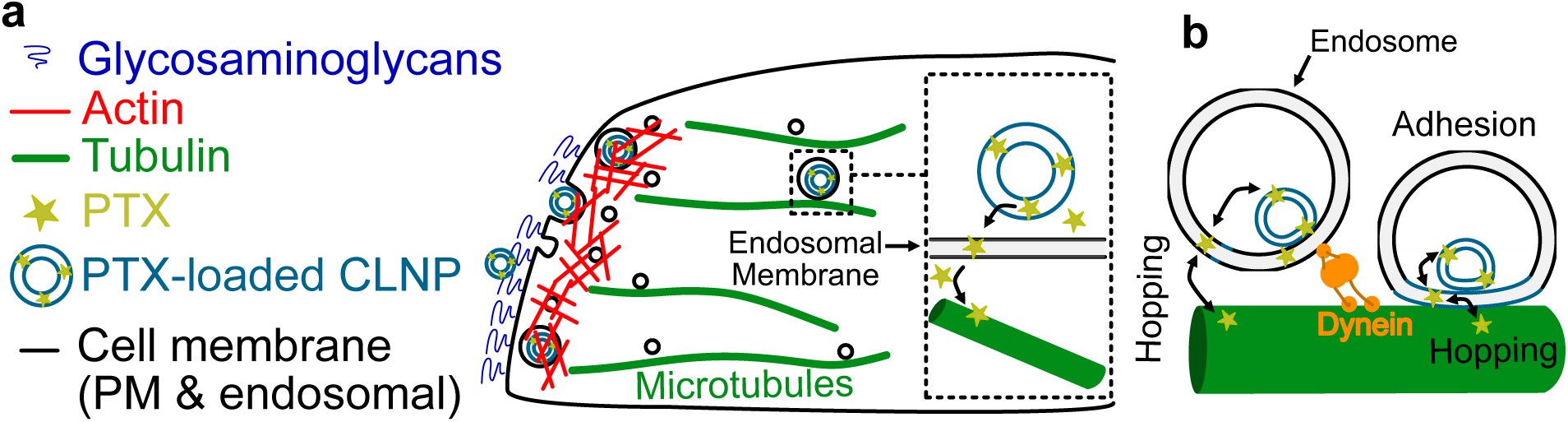
Graphic depicting the proposed mechanism of CLNP uptake and delivery of PTX to a cell. Upon endocytosis, endocytic vesicles containing CLNPs must diffuse across the ≈200 nm pore size actin mesh network near the cell surface before reaching microtubules and microtubule-associated motor proteins which can then rapidly transport those vesicles into the cell interior. Once brought into proximity with microtubules via endocytic vesicles, hydrophobic PTX molecules can hop from the CLNP membrane to their pharmacological target, the microtubules (**a**). CLNP loading in endocytic vesicles also contributes to the alteration of the effective charge of endocytic vesicles, both through attractive interactions between CLNPs and the endocytic vesicle membrane and cationic lipid hopping into the endocytic vesicle membrane, that may facilitate the adhesion of the endocytic vesicles containing CLNPs onto the microtubule (**b**). Hopping of PTX from endocytic vesicle membrane may occur both when the endocytic vesicle is tethered to a microtubule via a motor protein or when it adheres to the microtubule.

Figure 11 shows traces of cell borders obtained from DIC microscopy images that are overlayed on fluorescence microscopy images of *live* M21 cells treated with sonicated, 50 mol% MVL5 or DOTAP CLNPs containing fluorescent lipid label with and without 10 mol% PEG5k-DOPE. Images from the fluorescent lipid channel show that bare CLNPs form larger puncta, with DOTAP-CLNPs accumulating near the cell border while MVL5-CLNPs appear dispersed across the cell. In contrast, PEGylated MVL5- and DOTAP-CLNPs form small puncta which are nearly indistinguishable at the magnification used and are dispersed across the cell. This demonstrates that uptake of bare and PEGylated CLNPs follows a similar trend in live cells. The apparent dispersion of bare MVL5-CLNPs across cells is likely a result of insufficient resolution along the Z-plane to distinguish lipid puncta on top of cells from those in the cell body at this magnification versus the magnification used for Figure 8 and Figure 9.

**Figure 11.**
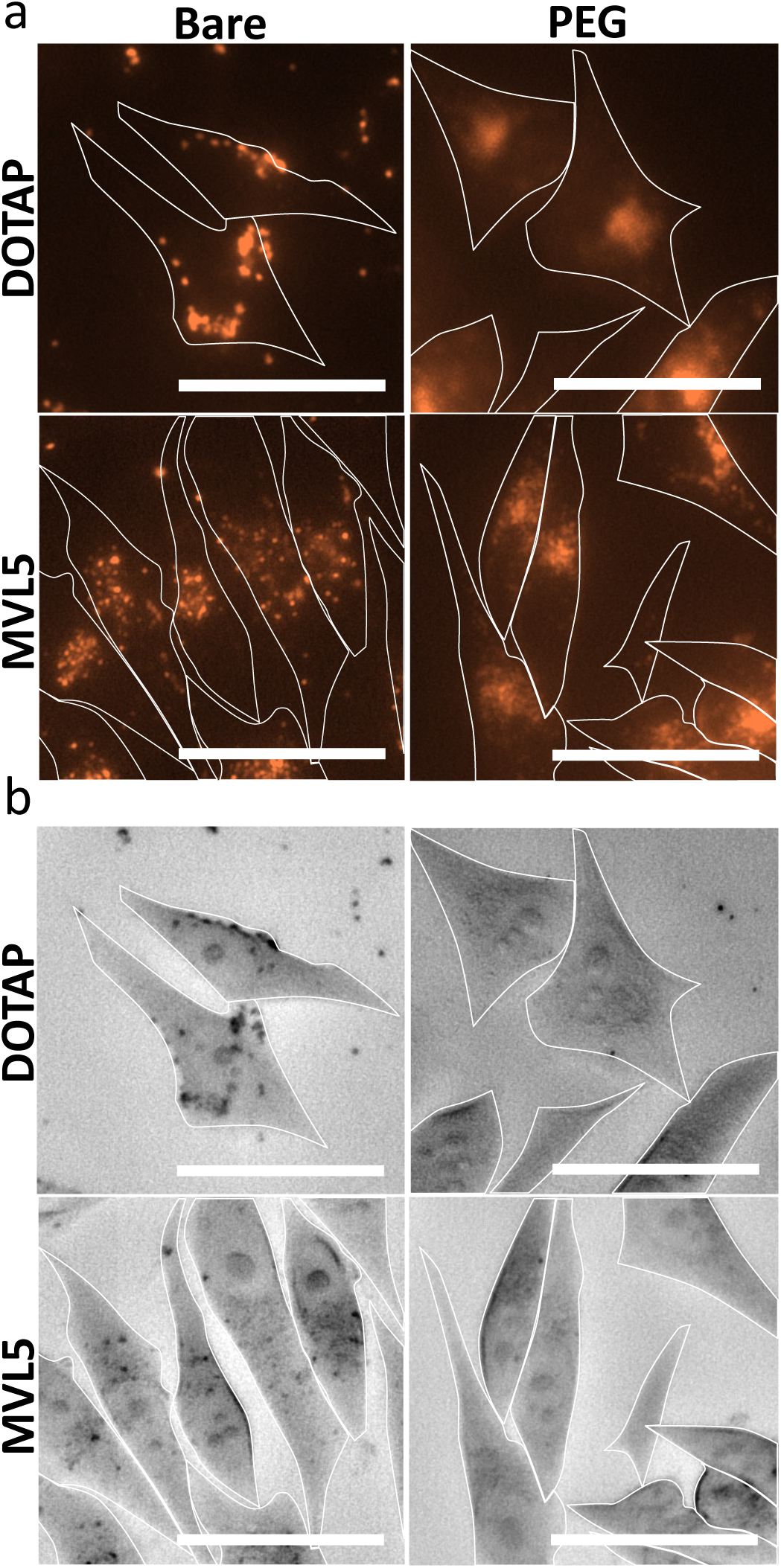
Fluorescent and DIC microscopy images of live M21 cells treated with fluorescently labeled bare and PEGylated CLNPs. M21 cells were treated with CLNPs of molar composition MVL5/DOPC/PEG-lipid/TRITC-DHPE/OG-PTX equal to 50/46.5−X_PEG_/X_PEG_/0.5/3 mol % or DOTAP/DOPC/PEG-lipid/TRITC-DHPE/OG-PTX equal to 50/46.5−X_PEG_/X_PEG_/0.5/3 mol % with either 0 or 10 mol % PEG-lipid. Fluorescent images of TRITC-DHPE signal in treated cells are shown in (**a**). The outlines of cells were determined using DIC images shown in (**b**) and are overlayed as white traces. Scale bars are 50mm.

## CONCLUSION

The multivalent lipid MVL5 is a high positive curvature cationic lipid that forms nanoparticles with disc, short rod, and spherical micelle structures when incorporated in DOPC lipid membranes. PEGylation further transitions rod micelles to elongated flexible rod. We investigated the capacity of these micellar MVL5 cationic lipid nanoparticles (CLNPs) to load the hydrophobic chemotherapeutic drug paclitaxel (PTX) and deliver PTX to human cancer cells. We found that MVL5 CLNPs have a PTX solubility limit that is nearly three times greater than DOTAP CLNPs based on the clinical EndoTAG-1^TM^ formulation. The improved PTX solubility of these MVL5 CLNPs translated into significantly improved cytotoxic efficacy for formulations at or above the PTX solubility limit for DOTAP, with PEGylation further enhancing this cytotoxic efficacy. Particle analysis of fluorescent microscopy images of cells treated with bare and PEGylated CLNPs revealed that PEGylation increases the uptake and penetration depth of MVL5 CLNPs into cells. The findings are consistent with a model where the rate limiting steps of effective PTX delivery include diffusion of sub-200 nm endocytic vesicles containing sterically stabilized PEGylated NPs across the actin mesh barrier to reach microtubules and hopping of PTX from NPs to endosomal vesicle membranes to microtubules, which are its pharmacological target. To further interrogate the accuracy of this model and resolve the factors most important for effective PTX delivery, future studies on the rate of PTX binding to microtubules when delivered to cells by bare versus PEGylated CLNPs, as well as the localization of PTX with CLNPs versus microtubules during delivery, are needed.

Our findings highlight the potential of PEGylated MVL5 CLNPs as superior therapeutic nanocarriers of PTX, and potentially numerous other hydrophobic drugs, to treat solid tumors and warrants the investigation of their tumor targeting and therapeutic performance *in vivo*. Such *in vivo* applications require carriers which stably incorporate drug over hours, preferentially target tumors through passive and ligand mediated means, and effectively enter tumor cells. The MVL5 CLNPs investigated in this study have significantly improved PTX solubility compared to CLs currently under clinical investigation and share the ability to leverage the EPR effect and electrostatic attraction to the highly negatively charged neoangiogenic vasculature associated with tumors. This is complemented by a sub 100 nm size and non-spherical shape profile which offers superior extravasation and cellular uptake potential when paired with steric stabilization from PEGylation. In addition, the ability to PEGylate these particles without compromising PTX solubility and cytotoxic efficacy means that ligand-mediated, cell-specific targeting is possible through conjugation of a targeting peptide onto a fraction of PEG moieties.^21, 60–62^ Finally, since the use of MVL5 in PEGylated CLNP carriers of PTX yielded significantly more effective PTX delivery, it is important that the PTX delivery efficacy of LNPs comprised of other multivalent lipids are explored to discover the properties of those lipids that might further enhance PTX delivery.

## Supporting Information

Cryo-TEM images of lipid nanoparticles containing mixtures of DOPC, DOTAP, or DOPC and PEG2K-lipid; DOPC and MVL5; and DOPC, PEG2K-lipid, and MVL5; wide field versions of fluorescent and DIC microscopy images of live M21 cells.

## Notes

The authors declare no competing financial interest.

## ACKNOWLEDGEMENTS

The authors dedicate the paper to the memory of the biophysicist Erich Sackmann. The work was supported, in part, by the US National Science Foundation, Division of Materials Research, under award DMR-1807327. W.S.F. is grateful for a Connie Frank Fellowship Award to students conducting biomedical research for human health at UC Santa Barbara.

## References

(1) Wani, M. C.; Taylor, H. L.; Wall, M. E.; Coggon, P.; McPhail, A. T. Plant antitumor agents. VI. Isolation and structure of taxol, a novel antileukemic and antitumor agent from Taxus brevifolia. Journal of the American Chemical Society 1971, 93 (9), 2325–2327. DOI: 10.1021/ja00738a045.

(2) Jordan, M. A.; Wilson, L. Microtubules as a target for anticancer drugs. Nature Reviews Cancer 2004, 4 (4), 253–265. DOI: 10.1038/nrc1317.

(3) Weaver, B. A. How Taxol/paclitaxel kills cancer cells. Mol Biol Cell 2014, 25 (18), 2677–2681. DOI: 10.1091/mbc.E14-04-0916 From NLM Medline.

(4) Rowinsky Eric, K.; Donehower Ross, C. Paclitaxel (Taxol). New Engl J Med 1995, 332 (15), 1004–1014. DOI: 10.1056/NEJM199504133321507 (acccessed 2025/02/24).

(5) Sofias, A. M.; Dunne, M.; Storm, G.; Allen, C. The battle of "nano" paclitaxel. Adv Drug Deliv Rev 2017, 122, 20–30. DOI: 10.1016/j.addr.2017.02.003 From NLM Medline.

(6) Dranitsaris, G.; Yu, B.; Wang, L.; Sun, W.; Zhou, Y.; King, J.; Kaura, S.; Zhang, A.; Yuan, P. Abraxane(R) versus Taxol(R) for patients with advanced breast cancer: A prospective time and motion analysis from a Chinese health care perspective. J Oncol Pharm Pract 2016, 22 (2), 205–211. DOI: 10.1177/1078155214556008 From NLM Medline.

(7) Gradishar, W. J.; Tjulandin, S.; Davidson, N.; Shaw, H.; Desai, N.; Bhar, P.; Hawkins, M.; O’Shaughnessy, J. Phase III Trial of Nanoparticle Albumin-Bound Paclitaxel Compared With Polyethylated Castor Oil–Based Paclitaxel in Women With Breast Cancer. Journal of Clinical Oncology 2016, 23 (31), 7794–7803. DOI: 10.1200/JCO.2005.04.937 (acccessed 2025/02/24).

(8) Rugo, H. S.; Barry, W. T.; Moreno-Aspitia, A.; Lyss, A. P.; Cirrincione, C.; Leung, E.; Mayer, E. L.; Naughton, M.; Toppmeyer, D.; Carey, L. A.;, et al. Randomized Phase III Trial of Paclitaxel Once Per Week Compared With Nanoparticle Albumin-Bound Nab-Paclitaxel Once Per Week or Ixabepilone With Bevacizumab As First-Line Chemotherapy for Locally Recurrent or Metastatic Breast Cancer: CALGB 40502/NCCTG N063H (Alliance). Journal of Clinical Oncology 2015, 33 (21), 2361–2369. DOI: 10.1200/JCO.2014.59.5298 (acccessed 2025/02/24).

(9) Weiss, R. B.; Donehower, R. C.; Wiernik, P. H.; Ohnuma, T.; Gralla, R. J.; Trump, D. L.; Baker, J. R.; Van Echo, D. A.; Von Hoff, D. D.; Leyland-Jones, B. Hypersensitivity reactions from taxol. Journal of Clinical Oncology 1990, 8 (7), 1263–1268. DOI: 10.1200/JCO.1990.8.7.1263 (acccessed 2025/02/24).

(10) Gelderblom, H.; Verweij, J.; Nooter, K.; Sparreboom, A. Cremophor EL: the drawbacks and advantages of vehicle selection for drug formulation. European Journal of Cancer 2001, 37 (13), 1590–1598. DOI: 10.1016/S0959-8049(01)00171-X.

(11) Dorr, R. T. Pharmacology and Toxicology of Cremophor EL Diluent. Annals of Pharmacotherapy 1994, 28 (5_suppl), S11-S14. DOI: 10.1177/10600280940280S503 (acccessed 2025/02/24).

(12) Safinya, C. R.; Ewert, K. K.; Majzoub, R. N.; Leal, C. Cationic liposome-nucleic acid complexes for gene delivery and gene silencing. New J Chem 2014, 38 (11), 5164–5172. DOI: 10.1039/C4NJ01314J From NLM PubMed-not-MEDLINE.

(13) Allen, T. M.; Cullis, P. R. Liposomal drug delivery systems: from concept to clinical applications. Adv Drug Deliv Rev 2013, 65 (1), 36–48. DOI: 10.1016/j.addr.2012.09.037 From NLM Medline.

(14) Hong, S. S.; Choi, J. Y.; Kim, J. O.; Lee, M. K.; Kim, S. H.; Lim, S. J. Development of paclitaxel-loaded liposomal nanocarrier stabilized by triglyceride incorporation. Int J Nanomedicine 2016, 11, 4465–4477. DOI: 10.2147/IJN.S113723 From NLM Publisher.

(15) Fetterly, G. J.; Straubinger, R. M. Pharmacokinetics of paclitaxel-containing liposomes in rats. AAPS PharmSci 2015, 5 (4), 32. DOI: 10.1208/ps050432.

(16) Sharma, A.; Sharma, U. S.; Straubinger, R. M. Paclitaxel-liposomes for intracavitary therapy of intraperitoneal P388 leukemia. Cancer Letters 1996, 107 (2), 265–272. DOI: 10.1016/0304-3835(96)04380-7.

(17) Zhou, R.; Mazurchuk, R. V.; Tamburlin, J. H.; Harrold, J. M.; Mager, D. E.; Straubinger, R. M. Differential pharmacodynamic effects of paclitaxel formulations in an intracranial rat brain tumor model. J Pharmacol Exp Ther 2010, 332 (2), 479–488. DOI: 10.1124/jpet.109.160044 From NLM Medline.

(18) Wang, H.; Cheng, G.; Du, Y.; Ye, L.; Chen, W.; Zhang, L.; Wang, T.; Tian, J.; Fu, F. Hypersensitivity reaction studies of a polyethoxylated castor oil-free, liposome-based alternative paclitaxel formulation. Mol Med Rep 2013, 7 (3), 947–952. DOI: 10.3892/mmr.2013.1264 From NLM Medline.

(19) Xu, X.; Wang, L.; Xu, H. Q.; Huang, X. E.; Qian, Y. D.; Xiang, J. Clinical comparison between paclitaxel liposome (Lipusu(R)) and paclitaxel for treatment of patients with metastatic gastric cancer. Asian Pac J Cancer Prev 2013, 14 (4), 2591–2594. DOI: 10.7314/apjcp.2013.14.4.2591 From NLM Medline.

(20) Zhang, J. A.; Anyarambhatla, G.; Ma, L.; Ugwu, S.; Xuan, T.; Sardone, T.; Ahmad, I. Development and characterization of a novel Cremophor EL free liposome-based paclitaxel (LEP-ETU) formulation. Eur J Pharm Biopharm 2005, 59 (1), 177–187. DOI: 10.1016/j.ejpb.2004.06.009 From NLM Medline.

(21) Koudelka, S.; Turanek, J. Liposomal paclitaxel formulations. J Control Release 2012, 163 (3), 322–334. DOI: 10.1016/j.jconrel.2012.09.006 From NLM Medline.

(22) Tahmasbi Rad, A.; Chen, C. W.; Aresh, W.; Xia, Y.; Lai, P. S.; Nieh, M. P. Combinational Effects of Active Targeting, Shape, and Enhanced Permeability and Retention for Cancer Theranostic Nanocarriers. ACS Appl Mater Interfaces 2019, 11 (11), 10505–10519. DOI: 10.1021/acsami.8b21609 From NLM Medline.

(23) Matsumura, Y.; Maeda, H. A new concept for macromolecular therapeutics in cancer chemotherapy: mechanism of tumoritropic accumulation of proteins and the antitumor agent smancs. Cancer Res 1986, 46 (12 Pt 1), 6387-6392, Research Support, Non-U.S. Gov’t.

(24) Schmitt-Sody, M.; Strieth, S.; Krasnici, S.; Sauer, B.; Schulze, B.; Teifel, M.; Michaelis, U.; Naujoks, K.; Dellian, M. Neovascular targeting therapy: paclitaxel encapsulated in cationic liposomes improves antitumoral efficacy. Clin Cancer Res 2003, 9 (6), 2335–2341, Research Support, Non-U.S. Gov’t.

(25) Thurston, G.; McLean, J. W.; Rizen, M.; Baluk, P.; Haskell, A.; Murphy, T. J.; Hanahan, D.; McDonald, D. M. Cationic liposomes target angiogenic endothelial cells in tumors and chronic inflammation in mice. J Clin Invest 1998, 101 (7), 1401–1413, Research Support, U.S. Gov’t, P.H.S. DOI: 10.1172/JCI965 [doi].

(26) Dellian, M.; Yuan, F.; Trubetskoy, V. S.; Torchilin, V. P.; Jain, R. K. Vascular permeability in a human tumour xenograft: molecular charge dependence. Br J Cancer 2000, 82 (9), 1513–1518, Research Support, Non-U.S. Gov’t Research Support, U.S. Gov’t, P.H.S. DOI: S0007092099911710 [pii] 10.1054/bjoc.1999.1171 [doi].

(27) Campbell, R. B.; Fukumura, D.; Brown, E. B.; Mazzola, L. M.; Izumi, Y.; Jain, R. K.; Torchilin, V. P.; Munn, L. L. Cationic charge determines the distribution of liposomes between the vascular and extravascular compartments of tumors. Cancer Res 2002, 62 (23), 6831–6836, Research Support, U.S. Gov’t, P.H.S.

(28) Steffes, V. M.; Murali, M. M.; Park, Y.; Fletcher, B. J.; Ewert, K. K.; Safinya, C. R. Distinct solubility and cytotoxicity regimes of paclitaxel-loaded cationic liposomes at low and high drug content revealed by kinetic phase behavior and cancer cell viability studies. Biomaterials 2017, 145, 242–255. DOI: 10.1016/j.biomaterials.2017.08.026 From NLM Medline.

(29) Sharma, A.; Straubinger, R. M. Novel taxol formulations: preparation and characterization of taxol-containing liposomes. Pharm Res 1994, 11 (6), 889–896, Research Support, U.S. Gov’t, P.H.S. DOI: 10.1023/a:1018994111594 [doi].

(30) Castro, J.; V. Tapia, L.; Silveyra, R.; Martinez Perez, C.; Deymier, P. Negative Impact of Paclitaxel Crystallization on Hydrogels and Novel Approaches for Anticancer Drug Delivery Systems. 2011.

(31) Que, C.; Gao, Y.; Raina, S. A.; Zhang, G. G. Z.; Taylor, L. S. Paclitaxel Crystal Seeds with Different Intrinsic Properties and Their Impact on Dissolution of Paclitaxel-HPMCAS Amorphous Solid Dispersions. Crystal Growth & Design 2018, 18 (3), 1548–1559. DOI: 10.1021/acs.cgd.7b01521.

(32) Zhen, Y.; Ewert, K. K.; Fisher, W. S.; Steffes, V. M.; Li, Y.; Safinya, C. R. Paclitaxel loading in cationic liposome vectors is enhanced by replacement of oleoyl with linoleoyl tails with distinct lipid shapes. Sci Rep 2021, 11 (1), 7311. DOI: 10.1038/s41598-021-86484-9 From NLM Medline.

(33) Allen, T. M.; Hansen, C. B.; Demenezes, D. E. L. Pharmacokinetics of Long-Circulating Liposomes. Adv Drug Deliver Rev 1995, 16 (2-3), 267–284. DOI: Doi 10.1016/0169- 409x(95)00029-7.

(34) Steffes, V. M.; Zhang, Z.; Ewert, K. K.; Safinya, C. R. Cryo-TEM Reveals the Influence of Multivalent Charge and PEGylation on Shape Transitions in Fluid Lipid Assemblies: From Vesicles to Discs, Rods, and Spheres. Langmuir 2023, 39 (50), 18424–18436. DOI: 10.1021/acs.langmuir.3c02664 From NLM Medline.

(35) Steffes, V. M.; Zhang, Z.; MacDonald, S.; Crowe, J.; Ewert, K. K.; Carragher, B.; Potter, C. S.; Safinya, C. R. PEGylation of Paclitaxel-Loaded Cationic Liposomes Drives Steric Stabilization of Bicelles and Vesicles thereby Enhancing Delivery and Cytotoxicity to Human Cancer Cells. ACS Appl Mater Interfaces 2020, 12 (1), 151–162. DOI: 10.1021/acsami.9b16150 From NLM Medline.

(36) Lipowsky, R. Bending of Membranes by Anchored Polymers. Europhys Lett 1995, 30 (4), 197–202. DOI: Doi 10.1209/0295-5075/30/4/002.

(37) Safran, S. Statistical Thermodynamics Of Surfaces, Interfaces, And Membranes; CRC Press, 2003. DOI: 10.1201/9780429497131.

(38) Aresh, W.; Liu, Y.; Sine, J.; Thayer, D.; Puri, A.; Huang, Y. K.; Wang, Y.; Nieh, M. P. The Morphology of Self-Assembled Lipid-Based Nanoparticles Affects Their Uptake by Cancer Cells. J Biomed Nanotechnol 2016, 12 (10), 1852–1863. DOI: 10.1166/jbn.2016.2292.

(39) Kinnear, C.; Moore, T. L.; Rodriguez-Lorenzo, L.; Rothen-Rutishauser, B.; Petri-Fink, A. Form Follows Function: Nanoparticle Shape and Its Implications for Nanomedicine. Chem Rev 2017, 117 (17), 11476–11521. DOI: 10.1021/acs.chemrev.7b00194.

(40) Simon-Gracia, L.; Scodeller, P.; Fisher, W. S.; Sidorenko, V.; Steffes, V. M.; Ewert, K. K.; Safinya, C. R.; Teesalu, T. Paclitaxel-Loaded Cationic Fluid Lipid Nanodiscs and Liposomes with Brush-Conformation PEG Chains Penetrate Breast Tumors and Trigger Caspase-3 Activation. ACS Appl Mater Interfaces 2022, 14 (51), 56613–56622. DOI: 10.1021/acsami.2c17961 From NLM Medline.

(41) Ewert, K.; Ahmad, A.; Evans, H. M.; Schmidt, H. W.; Safinya, C. R. Efficient synthesis and cell-transfection properties of a new multivalent cationic lipid for nonviral gene delivery. J Med Chem 2002, 45 (23), 5023–5029. DOI: 10.1021/jm020233w.

(42) Farago, O.; Ewert, K.; Ahmad, A.; Evans, H. M.; Gronbech-Jensen, N.; Safinya, C. R. Transitions between distinct compaction regimes in complexes of multivalent cationic lipids and DNA. Biophysical Journal 2008, 95 (2), 836–846. DOI: 10.1529/biophysj.107.124669.

(43) Zidovska, A.; Ewert, K. K.; Quispe, J.; Carragher, B.; Potter, C. S.; Safinya, C. R. Block liposome and nanotube formation is a general phenomenon of two-component membranes containing multivalent lipids. Soft Matter 2011, 7 (18), 8363–8369. DOI: 10.1039/c1sm05481c.

(44) Leal, C.; Ewert, K. K.; Shirazi, R. S.; Bouxsein, N. F.; Safinya, C. R. Nanogyroids Incorporating Multivalent Lipids: Enhanced Membrane Charge Density and Pore Forming Ability for Gene Silencing. Langmuir 2011, 27 (12), 7691–7697. DOI: 10.1021/la200679x.

(45) Bouxsein, N. F.; McAllister, C. S.; Ewert, K. K.; Samuel, C. E.; Safinya, C. R. Structure and gene silencing activities of monovalent and pentavalent cationic lipid vectors complexed with siRNA. Biochemistry 2007, 46 (16), 4785–4792. DOI: 10.1021/bi062138l.

(46) Ahmad, A.; Evans, H. M.; Ewert, K.; George, C. X.; Samuel, C. E.; Safinya, C. R. New multivalent cationic lipids reveal bell curve for transfection efficiency versus membrane charge density: lipid-DNA complexes for gene delivery. J Gene Med 2005, 7 (6), 739–748. DOI: 10.1002/jgm.717.

(47) Pollard, T. D. E., W. C.; Lippincott-Schwartz, J.; Johnson, G. T. Cell Biology; Elsevier, 2017.

(48) Campbell, R. B.; Balasubramanian, S. V.; Straubinger, R. M. Influence of cationic lipids on the stability and membrane properties of paclitaxel-containing liposomes. J Pharm Sci 2001, 90 (8), 1091–1105. DOI: 10.1002/jps.1063 From NLM Medline.

(49) Kannan, V.; Balabathula, P.; Divi, M. K.; Thoma, L. A.; Wood, G. C. Optimization of drug loading to improve physical stability of paclitaxel-loaded long-circulating liposomes. J Liposome Res 2015, 25 (4), 308–315. DOI: 10.3109/08982104.2014.995671.

(50) Bernsdorff, C.; Reszka, R.; Winter, R. Interaction of the anticancer agent Taxol™ (paclitaxel) with phospholipid bilayers. J Biomed Mater Res 1999, 46 (2), 141–149. DOI: Doi 10.1002/(Sici)1097-4636(199908)46:2<141::Aid-Jbm2>3.0.Co;2-U.

(51) Johnsson, M.; Edwards, K. Liposomes, disks, and spherical micelles: Aggregate structure in mixtures of gel phase phosphatidylcholines and poly(ethylene glycol)-phospholipids. Biophysical Journal 2003, 85 (6), 3839–3847. DOI: Doi 10.1016/S0006-3495(03)74798-5.

(52) Rad, A. T.; Chen, C. W.; Aresh, W.; Xia, Y.; Lai, P. S.; Nieh, M. P. Combinational Effects of Active Targeting, Shape, and Enhanced Permeability and Retention for Cancer Theranostic Nanocarriers. Acs Appl Mater Inter 2019, 11 (11), 10505–10519. DOI: 10.1021/acsami.8b21609.

(53) Rad, A. T.; Hargrove, D.; Daneshmandi, L.; Ramsdell, A.; Lu, X. L.; Nieh, M. P. Codelivery of Paclitaxel and Parthenolide in Discoidal Bicelles for a Synergistic Anticancer Effect: Structure Matters. Adv Nanobiomed Res 2022, 2 (1). DOI: ARTN 2100080 10.1002/anbr.202100080.

(54) Steffes, V.; MacDonald, S.; Crowe, J.; Murali, M.; Ewert, K. K.; Li, Y.; Safinya, C. R. Lipids with negative spontaneous curvature decrease the solubility of the cancer drug paclitaxel in liposomes. Eur Phys J E Soft Matter 2023, 46 (12), 128. DOI: 10.1140/epje/s10189-023-00388-2 From NLM Medline.

(55) Eichhorn, M. E.; Ischenko, I.; Luedemann, S.; Strieth, S.; Papyan, A.; Werner, A.; Bohnenkamp, H.; Guenzi, E.; Preissler, G.; Michaelis, U.;, et al. Vascular targeting by EndoTAG- 1 enhances therapeutic efficacy of conventional chemotherapy in lung and pancreatic cancer. Int J Cancer 2010, 126 (5), 1235–1245. DOI: 10.1002/ijc.24846 From NLM Medline.

(56) Eichhorn, M. E.; Luedemann, S.; Strieth, S.; Papyan, A.; Ruhstorfer, H.; Haas, H.; Michaelis, U.; Sauer, B.; Teifel, M.; Enders, G.;, et al. Cationic lipid complexed camptothecin (EndoTAG-2) improves antitumoral efficacy by tumor vascular targeting. Cancer Biol Ther 2007, 6 (6), 920–929. DOI: 10.4161/cbt.6.6.4207 From NLM Medline.

(57) Svitkina, T. M. Actin Cell Cortex: Structure and Molecular Organization. Trends Cell Biol 2020, 30 (7), 556–565. DOI: 10.1016/j.tcb.2020.03.005 From NLM Medline.

(58) Chugh, P.; Paluch, E. K. The actin cortex at a glance. J Cell Sci 2018, 131 (14). DOI: 10.1242/jcs.186254 From NLM Medline.

(59) Raviv, U.; Needleman, D. J.; Li, Y. L.; Miller, H. P.; Wilson, L.; Safinya, C. R. Cationic liposome-microtubule complexes: Pathways to the formation of two-state lipid-protein nanotubes with open or closed ends. P Natl Acad Sci USA 2005, 102 (32), 11167–11172. DOI: 10.1073/pnas.0502183102.

(60) Ewert, K. K.; Kotamraju, V. R.; Majzoub, R. N.; Steffes, V. M.; Wonder, E. A.; Teesalu, T.; Ruoslahti, E.; Safinya, C. R. Synthesis of linear and cyclic peptide-PEG-lipids for stabilization and targeting of cationic liposome-DNA complexes. Bioorg Med Chem Lett 2016, 26 (6), 1618–1623. DOI: 10.1016/j.bmcl.2016.01.079 From NLM Medline.

(61) Biswas, S.; Dodwadkar, N. S.; Deshpande, P. P.; Torchilin, V. P. Liposomes loaded with paclitaxel and modified with novel triphenylphosphonium-PEG-PE conjugate possess low toxicity, target mitochondria and demonstrate enhanced antitumor effects and. Journal of Controlled Release 2012, 159 (3), 393–402. DOI: 10.1016/j.jconrel.2012.01.009.

(62) MacEwan, S. R.; Chilkoti, A. Harnessing the power of cell-penetrating peptides: activatable carriers for targeting systemic delivery of cancer therapeutics and imaging agents. Wires Nanomed Nanobi 2013, 5 (1), 31–48. DOI: 10.1002/wnan.1197.

