## Supplemental Information for "Multivalent Lipid MVL5 Micellar Nanoparticles Exhibit Dramatically Increased Loading of Paclitaxel with PEGylation Enhancing Human Cancer Cell Penetration Depth and Cytotoxicity"

### Contents

Cryo-TEM Images of Lipid Nanoparticles Containing DOPC, DOTAP, or DOPC and PEG2K-lipid S1

Cryo-TEM Images of Cationic Lipid Nanoparticles Containing DOPC and Increasing Levels of MVL5 Content S2

Cryo-TEM Images of Lipid Nanoparticles Containing DOPC, PEG2K-lipid, and Increasing Levels of MVL5 Content S3

Fluorescent and DIC Microscopy Images of M21 Cells Treated with Fluorescently Labeled Bare DOTAP CLNPs S4

Fluorescent and DIC Microscopy Images of M21 Cells Treated with Fluorescently Labeled PEGylated DOTAP CLNPs S5

Fluorescent and DIC Microscopy Images of M21 Cells Treated with Fluorescently Labeled Bare MVL5 CLNPs S6

Fluorescent and DIC Microscopy Images of M21 Cells Treated with Fluorescently Labeled PEGylated MVL5 CLNPs S7

Wide Field Fluorescent and DIC Microscopy Images of Live M21 Cells Treated with DOTAP and MVL5 CLNPs with and without PEG-lipid Used for Figure 11 S8

### Cryo-TEM Images of Cationic Lipid Nanoparticles Containing DOPC, DOTAP, or DOPC and PEG2K-lipid.

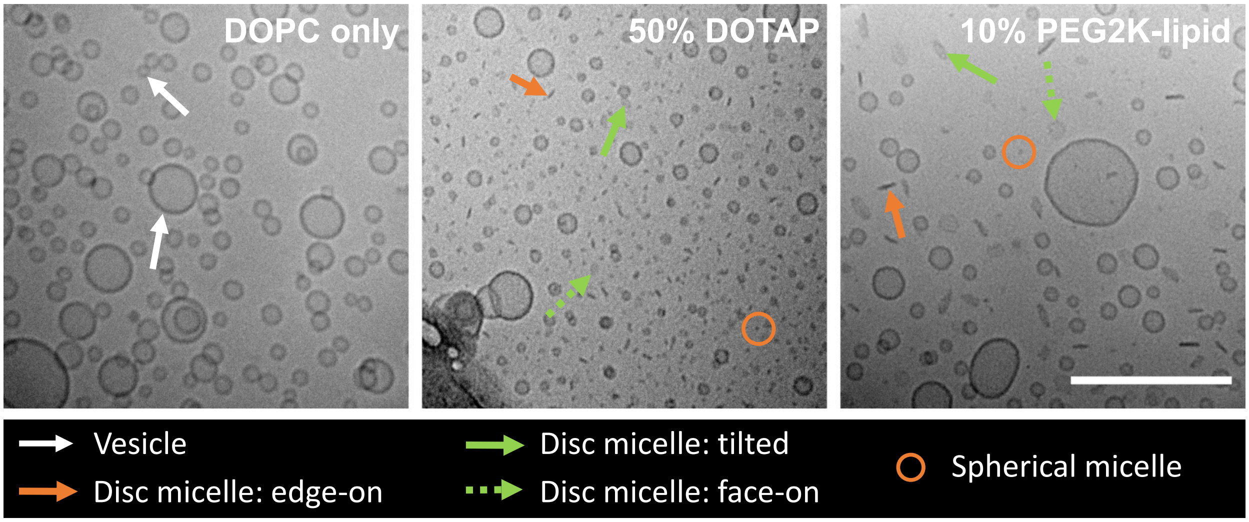


**Supplemental Figure 1. Addition of poly(ethylene glycol)2K-lipid (PEG2K-lipid) or univalent cationic lipid 2,3-dioleyloxypropyltrimethylammonium chloride (DOTAP, +1e) to neutral liposomes modifies their morphology.** Cryo-TEM showing morphologies in formulations containing charge-neutral lipid 1,2-dioleoyl-*sn*-glycero-phosphocholine (DOPC) (left panel), DOPC and 50 mol% DOTAP (middle panel), and DOPC and 10 mol% PEG2K-lipid (right panel). Arrows and circles point to typical vesicle and micelle structures as described in the legend. The majority of the non-vesicular objects are disc shaped micelles (middle and right panels), seen in tilted (green arrow), face-on (green dashed arrow) and edge-on (red arrow) orientation. Scale bar: 200 nm. The formulations contained 3 mol% of the hydrophobic drug paclitaxel (PTX), which does not alter morphologies. Images adapted with permission from reference.[34] Copyright 2020 American Chemical Society.

**Cryo-TEM Images of Cationic Lipid Nanoparticles Containing DOPC and Increasing Levels of MVL5 Content.**

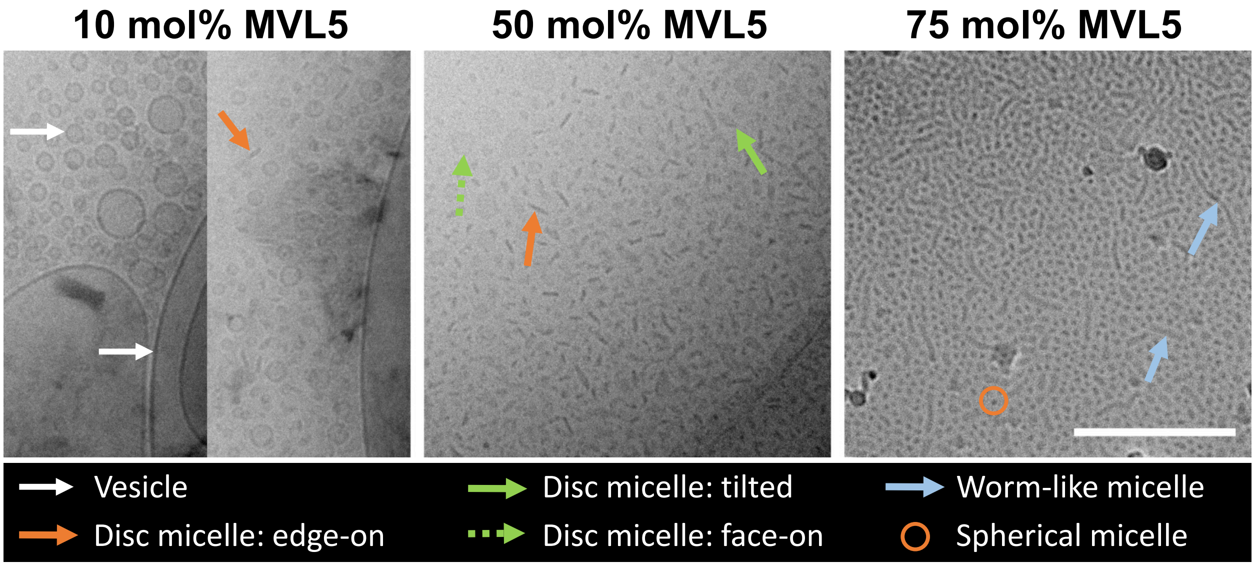


**Supplemental Figure 2. Liposomal morphologies of charge-neutral lipid 1,2-dioleoyl-*sn*-glycero-phosphocholine (DOPC) and multivalent lipid MVL5 (+5e) formulations at different concentrations of MVL5 as specified above each panel.** Cryo-TEM images reveal that the incorporation of increasing fractions of the high curvature cationic lipid MVL5 changes the liposome morphology from vesicles (left panel) to discs (50 mol% MVL%, middle panel), to short/intermediate length worm-like micellar rods coexisting with spheres (75 mol% MVL5, right panel). Arrows and circles identify vesicle and micelle structures according to the legend. The formulation with 10 mol% MVL5 only showed presence of discs coexisting with vesicles in a single micrograph out of 38 micrographs (right half of left panel), with all other micrographs showing spherical vesicles. Scale bar: 200 nm. Images adapted with permission from ref.[34] Copyright 2020 American Chemical Society.

**Cryo-TEM Images of Lipid Nanoparticles Containing DOPC, PEG2K-lipid, and Increasing Levels of MVL5 Content.**

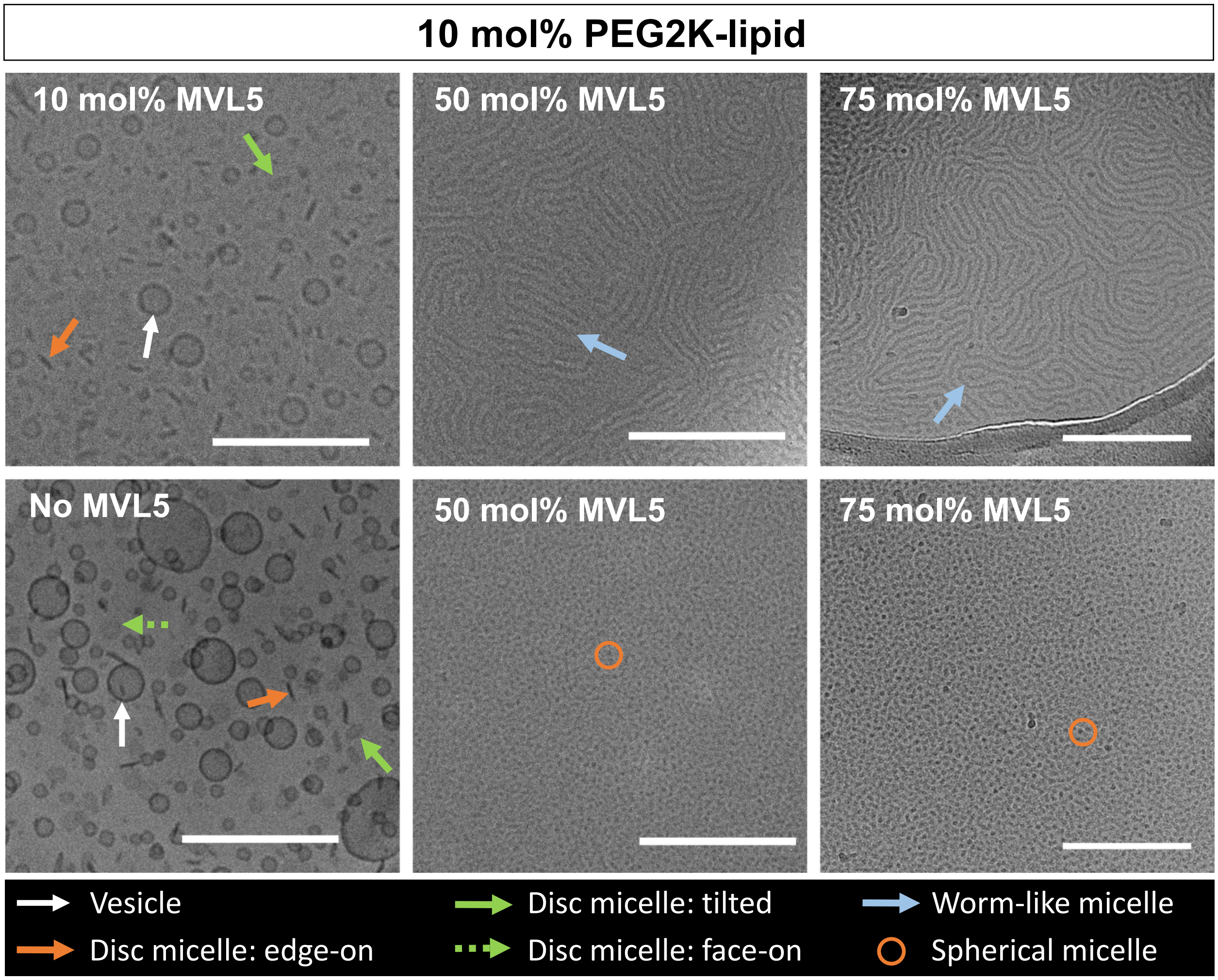


**Supplemental Figure 3. Morphologies of liposomal formulations in ternary mixtures consisting of multivalent lipid MVL5 (+5e), poly(ethylene glycol)2K-lipid (PEG2K-lipid), and charge-neutral lipid 1,2-dioleoyl-*sn*-glycero-phosphocholine (DOPC) revealed by CryoTEM (bottom left pane is a binary mixture of 10 mol% PEG2K-lipid and DOPC).** The left panels reveal coexistence of vesicles and disc micelles for 10 mol% PEG2K-lipid formulations with (top left panel) and without (bottom left panel) 10 mol% MVL. In formulations containing 10 mol% PEG2K-lipid and 50 mol% MVL5, flexible worm-like cylindrical micelles (top middle panel) coexist with spherical micelles (bottom middle panel). At 75 mol% MVL5 and 10 mol% PEG2K-lipid, spherical micelles (bottom right panel) are more abundant and coexist with very long and flexible cylindrical micelles (top right panel). Arrows and circles identify vesicle and micelle structures according to the legend. Lipid composition: 10 mol% PEG2K-lipid, MVL5 content as specified on the images, remainder DOPC. Scale bars: 200 nm. Images adapted with permission from reference.[34] Copyright 2020 American Chemical Society.

### Fluorescent and DIC Microscopy Images of M21 Cells Treated with Fluorescently Labeled Bare DOTAP CLNPs

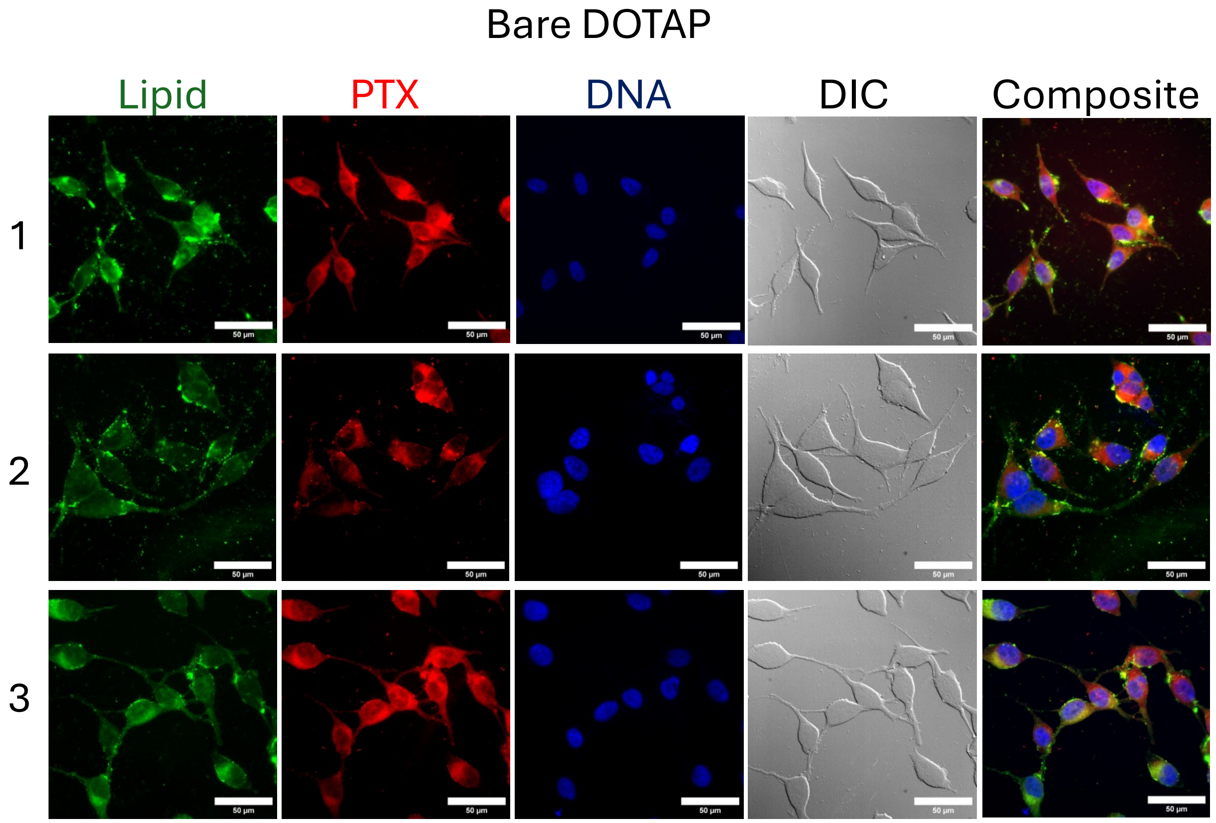


**Supplemental Figure 4. Selected z-slices from fluorescent microscopy image stacks of fixed M21 cells treated with bare DOTAP CLNPs containing fluorescently labeled lipid and PTX-fluorophore conjugate.** Images of M21 cells treated with CLNPs of molar composition DOTAP/DOPC/TRITC-DHPEE/Janelia-PTX equal to 50/46.5/0.5/3 mol% from three repeat experiments. CL lipid (TRITC-DHPE) signal is shown in green, PTX (Janelia-PTX) signal is shown in red, DAPI was used to stain cell nuclei after fixation and is shown in blue. Scale bars are 50 μm. These were used to quantify CLNP puncta number and penetration depth in cell profiler as described in the manuscript.

**Fluorescent and DIC Microscopy Images of M21 Cells Treated with Fluorescently Labeled PEGylated DOTAP CLNPs**

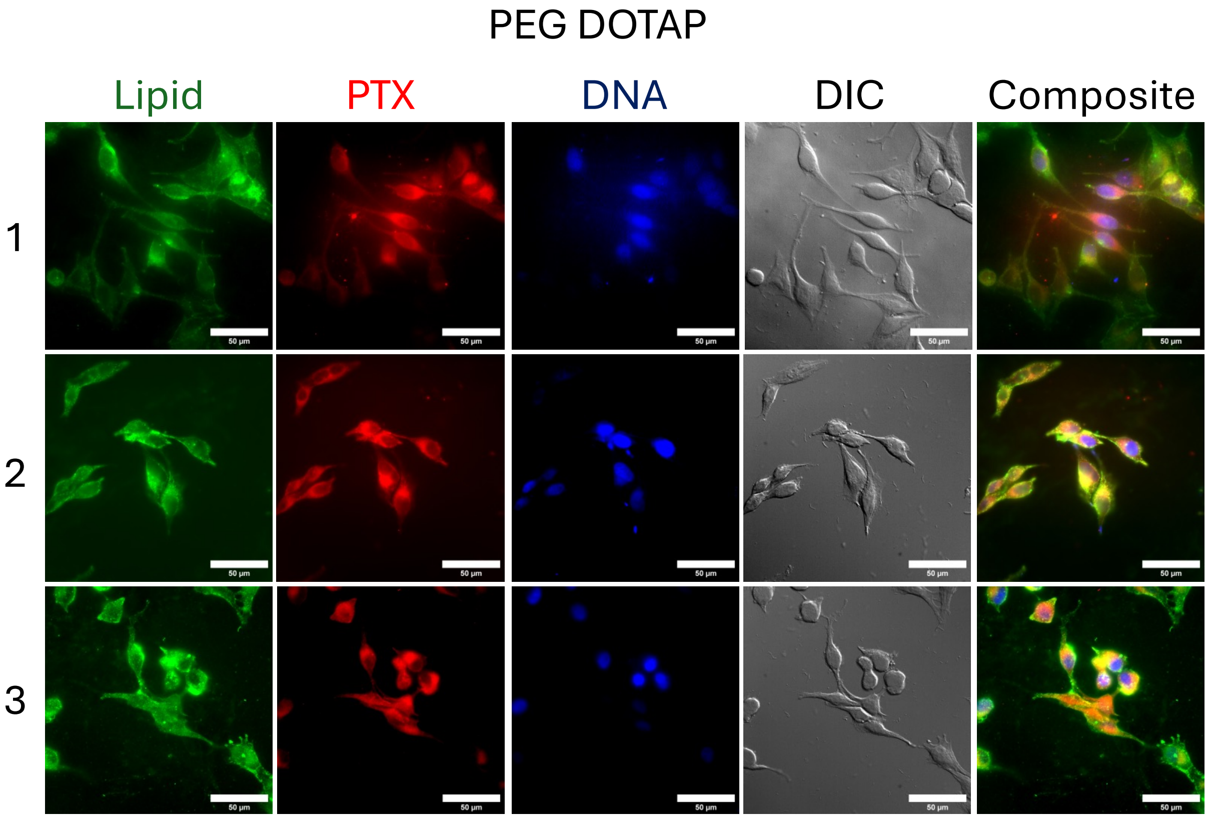


**Supplemental Figure 5. Selected z-slices from fluorescent microscopy image stacks of fixed M21 cells treated with PEGylated DOTAP CLNPs containing fluorescently labeled lipid and PTX-fluorophore conjugate.** Images of M21 cells treated with CLNPs of molar composition DOTAP/DOPC/PEG-lipid/TRITC-DHPEE/Janelia-PTX equal to 50/36.5/10/0.5/3 mol% from three repeat experiments. CL lipid (TRITC-DHPE) signal is shown in green, PTX (Janelia-PTX) signal is shown in red, DAPI was used to stain cell nuclei after fixation and is shown in blue. Scale bars are 50 μm. These were used to quantify CLNP puncta number and penetration depth in cell profiler as described in the manuscript.

**Fluorescent and DIC Microscopy Images of M21 Cells Treated with Fluorescently Labeled Bare MVL5 CLNPs**

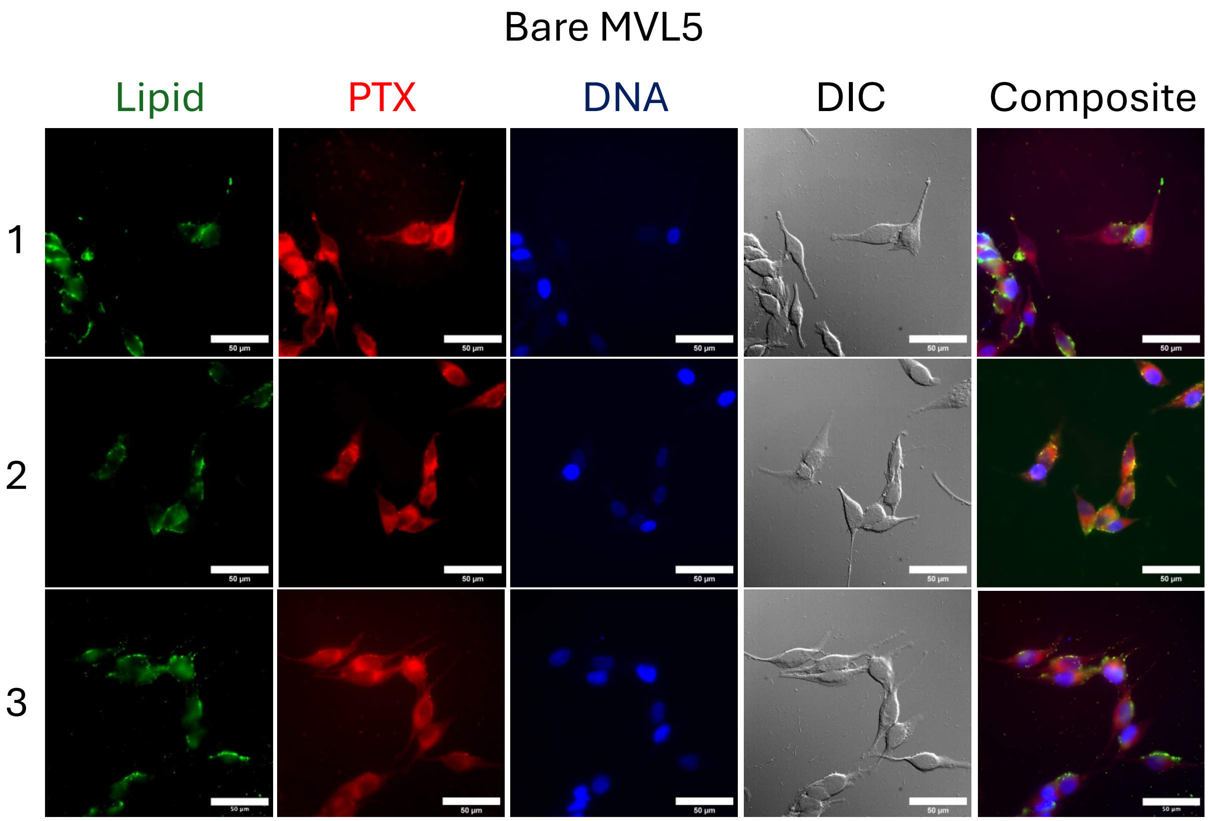


**Supplemental Figure 6. Selected z-slices from fluorescent microscopy image stacks of fixed M21 cells treated with bare MVL5 CLNPs containing fluorescently labeled lipid and PTX-fluorophore conjugate.** Images of M21 cells treated with CLNPs of molar composition MVL5/DOPC/TRITC-DHPEE/Janelia-PTX equal to 50/46.5/0.5/3 mol% from three repeat experiments. CL lipid (TRITC-DHPE) signal is shown in green, PTX (Janelia-PTX) signal is shown in red, DAPI was used to stain cell nuclei after fixation and is shown in blue. Scale bars are 50 µm. These were used to quantify CLNP puncta number and penetration depth in cell profiler as described in the manuscript.

**Fluorescent and DIC Microscopy Images of M21 Cells Treated with Fluorescently Labeled PEGylated MVL5 CLNPs**

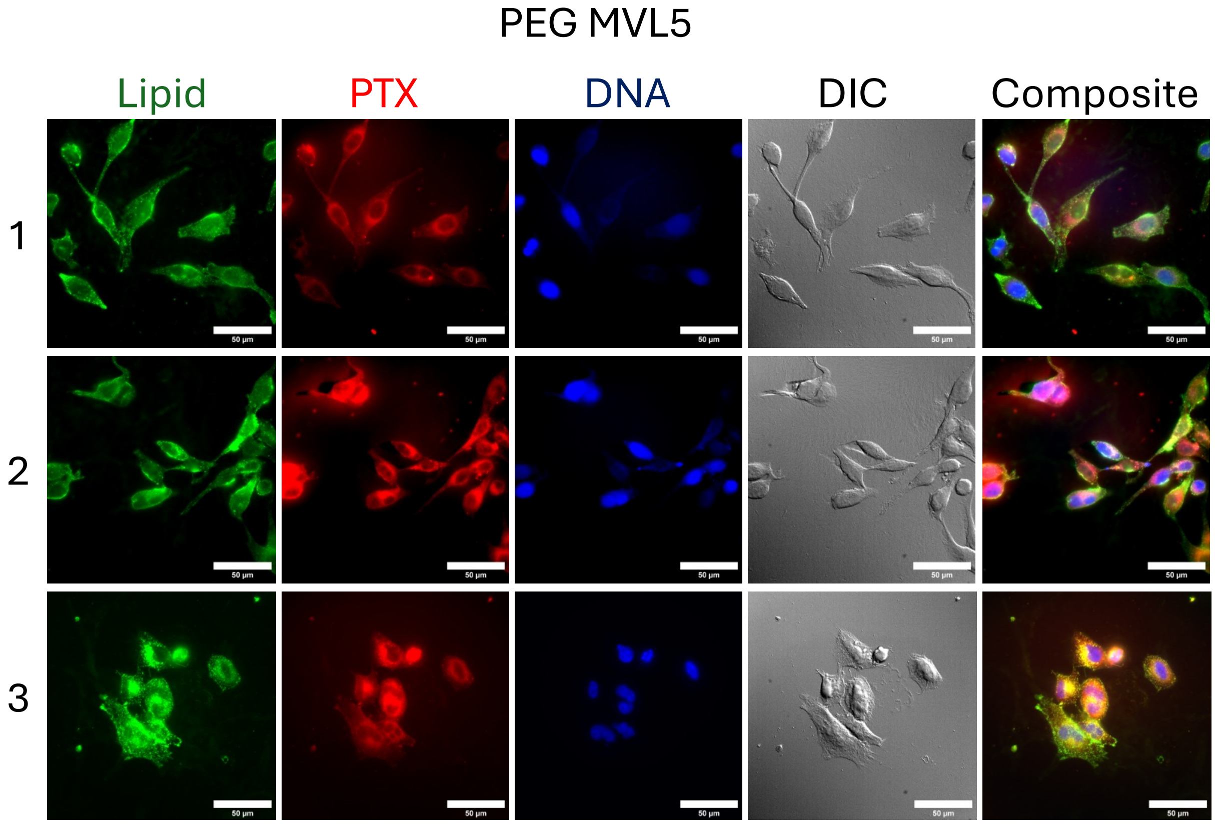


**Supplemental Figure 7. Selected z-slices from fluorescent microscopy image stacks of fixed M21 cells treated with PEGylated MVL5 CLNPs containing fluorescently labeled lipid and PTX-fluorophore conjugate.** Images of M21 cells treated with CLNPs of molar composition MVL5/DOPC/PEG-lipid/TRITC-DHPEE/Janelia-PTX equal to 50/36.5/10/0.5/3 mol% from three repeat experiments. CL lipid (TRITC-DHPE) signal is shown in green, PTX (Janelia-PTX) signal is shown in red, DAPI was used to stain cell nuclei after fixation and is shown in blue. Scale bars are 50 µm. These were used to quantify CLNP puncta number and penetration depth in cell profiler as described in the manuscript.

**Wide Field Fluorescent and DIC Microscopy Images of Live M21 Cells Treated with DOTAP and MVL5 CLNPs with and without PEG-lipid Used for Figure 11.**

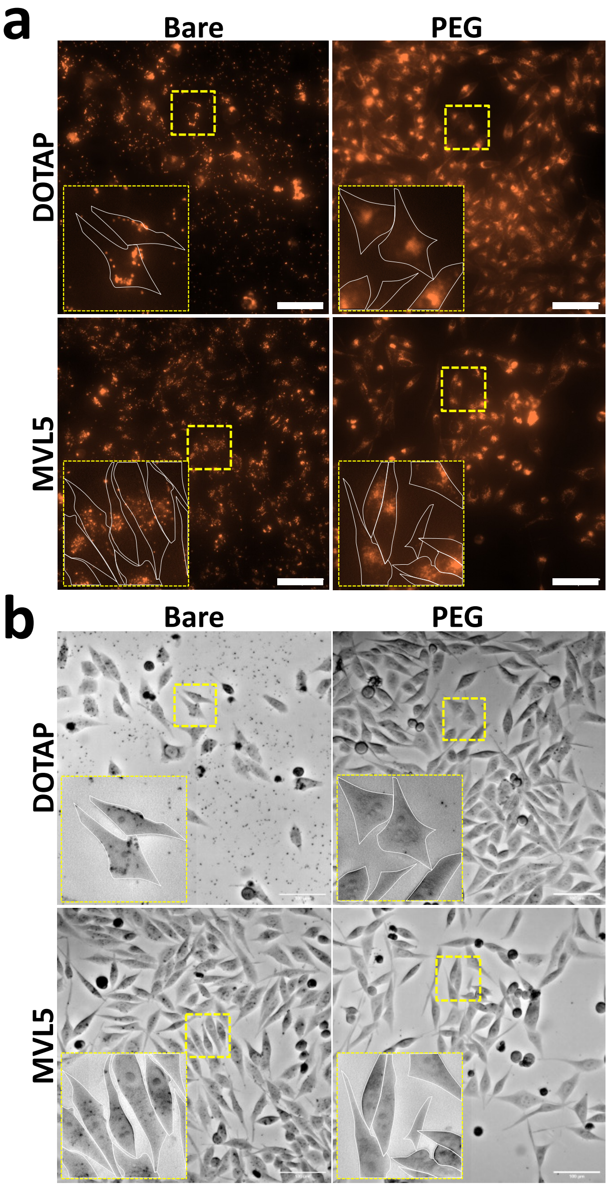


**Supplemental Figure 8**. **Wide field fluorescence and DIC images containing select regions used for live cell images in Figure 11.** Fluorescent and DIC microscopy images of live M21 cells treated with fluorescently labeled bare and PEGylated CLNPs. M21 cells were treated with CLNPs of molar composition MVL5/DOPC/PEG-lipid/TRITC-DHPE/OG-PTX equal to 50/46.5−$X_{PEG}$/$X_{PEG}$/0.5/3 mol % or DOTAP/DOPC/PEG-lipid/TRITC-DHPE/OG-PTX equal to 50/46.5−$X_{PEG}$/$X_{PEG}$/0.5/3 mol % with either 0 or 10 mol % PEG-lipid. Images show fluorescent signal emitted from the TRITC-DHPE probe incorporated in the CLNP membrane (b) or the outline of cells from DIC microscopy (b). Regions highlighted with dashed yellow squares are magnified in insets, with the outline of cells (determined using DIC images) overlayed in white traces. Scale bars are 100μm.
